# Light-up split Broccoli aptamer as a versatile tool for RNA assembly monitoring in cell-free TX-TL system, hybrid RNA/DNA origami tagging and DNA biosensing

**DOI:** 10.1101/2022.07.20.500791

**Authors:** Emanuela Torelli, Benjamin Shirt-Ediss, Silvia A. Navarro, Marisa Manzano, Priya Vizzini, Natalio Krasnogor

**Affiliations:** Interdisciplinary Computing and Complex BioSystems (ICOS), Centre for Synthetic Biology and Bioeconomy (CSBB), Newcastle University, United Kingdom; Dipartimento di Scienze AgroAlimentari, Ambientali e Animali, Università degli Studi di Udine, Italy

**Keywords:** Light-up split Broccoli aptamer, nucleic acid-based devices, *Campylobacter* spp., hybrid RNA/DNA origami, cell-free TX-TL system, therapeutic nanostructures monitoring

## Abstract

Binary light-up aptamers are intriguing and emerging tools with potential in different fields. Herein, we demonstrate the versatility of a split Broccoli aptamer system able to turn on the fluorescence signal only in the presence of a complementary sequence.

First, an RNA three-way junction harbouring the split system was assembled in an *E. coli* based cell-free TX-TL system where the folding of the functional aptamer is demonstrated. Then, the same strategy is introduced into a ‘bio-orthogonal’ hybrid RNA/DNA rectangle origami characterized by atomic force microscopy: the activation of the split system through the origami self-assembly is demonstrated. Finally, our system is successfully used to detect femtomoles of a *Campylobacter* spp. DNA target sequence.

Potential applications of our system include real-time monitoring of the self-assembly of nucleic acid-based devices *in vivo* and of intracellular delivery of therapeutic nanostructures, as well as *in vitro* and *in vivo* detection of different DNA/RNA target.

## INTRODUCTION

Light-up RNA aptamers offer an alternative to fluorescent probes and proteins (e.g., green fluorescent protein GFP, etc.), avoiding the covalent conjugation with fluorescent molecules and the alteration of natural proteins expression level and localization *in vivo*.^1^ The first GFP-mimicking light-up aptamer showed a response upon binding to malachite green, a highly cytotoxic dye.^2,3^ To develop new tools for RNA labelling and tracking in living cells, RNA aptamers able to bind a non-cytotoxic, cell-permeable and brighter dye (DFHBI and analogues) were successfully introduced and selected.^4-6^ To further improve RNA imaging and aptamer stability in a cellular environment, Filonov and coll.^7^ obtained a new light-up Broccoli aptamer derived from a combined SELEX-fluorescence activated cell sorting (FACS) approach: the library selected by SELEX was cloned into expression plasmids and transcribed in *Escherichia coli* cells. Expressed in both, prokaryotic and eukaryotic cells, Broccoli showed a high folding efficiency, a lower magnesium chloride dependence and an increased thermostability compared to Spinach and Spinach2 aptamers.^8^ Because of this, Broccoli aptamer, either as full or split sequences, received increased attention and found a variety of applications.^1^

The split Broccoli concept was firstly introduced *in vivo* by dividing the full sequence into two autonomous RNA strands, called Top and Bottom: the two-strand system was genetically encoded and able to assemble *Escherichia coli*.^9^ However, the two aptameric fragments could not hybridize to a selected target complementary sequence, which can represent a limitation when self-assembly reaction monitoring is considered. Most recently, split versions of the Broccoli aptamer were used *in vivo* as a reporter integrated into a catalytic hairpin assembly circuit,^10^ an aptamer-initiated fluorescence complementation assay^11^ and an *in situ* amplification method.^12^

In detail, the split system was integrated into a catalytic hairpin assembly circuit: the target triggered and catalysed the hybridization between two hairpins (H1 and H2) modified with split Broccoli, called Broc and Coli. However, once the aptamer was reconstituted, the target was disconnected,^10^ making the system unsuitable for stable nucleic acid self-assemblies (e.g., origami nanostructures). Following a different approach, the Broccoli reporter was combined with an RNA-based hybridization chain reaction for imaging RNA location: in the presence of an initiator RNA, H1 and H2 hairpins hybridized, resulting in a double-stranded concatameric assembly able to activate a fluorescence signal,^12^ but potentially disturb the cellular expression pattern. On the other hand, considering a simpler design developed to image native mRNA in mammalian cells, Wang and coll.^11^ used two RNA split fragments extended with recognition sequences at the UUCG loop. Compared to the previous approach, this imaging strategy minimally disturbed the natural behaviour of mRNA, the cell morphology and the cell viability.^11^ However, the system was not tested in prokaryotic cells, in which different transcribed terminators (e.g. a 47 nt T7 terminator instead of a 63 nt mini polyA terminator required by RNA pol II^11^) could result in different transcript composition and size, influencing a specific nucleic acid assembly. When we considered the split system^11^ as a component of a three-way junction with T7 terminators, the full Broccoli aptamer reconstitution was not successfully predicted.

Therefore, the described approaches^9,10,11,12^ are unsuitable when *in vivo* synthesis monitoring of stable DNA-encoded RNA origami^14^ and memory structures are considered as potentially generic interface to bacterial cell processes (e.g. transcription, translation, transduction, etc. events). For this reason and as a further step along the RNA origami synthesis^13,14^ in bacteria, here we first explore the use of our binary Broccoli aptamer^13,14^ to monitor the RNA self-assembly in an *E. coli* transcription-translation (TX-TL) system, which represents an intermediate step before moving encoded RNA assemblies to synthetic cells or bacterial cells. In detail, the split sequences, ending with 8 nt of a stabilizing bio-orthogonal F30 scaffold,^8^ were elongated in 5’ and 3’, while the 4 nt loop UUCG was removed from the stem.^13,14^ The designed RNA three-way junction system including the split sequences was genetically encoded and expressed in an *E. coli* cell-free system at 37 °C: the self-assembly and the fluorescence emission signal were successfully simulated, visualized and detected.

Furthermore, we demonstrate the versatility and adaptability of the binary system combining the split functionality with a rectangle RNA/DNA hybrid origami nanostructure and suggesting a biocompatible alternative to organic dyes for *in vitro* and *in vivo* applications. A ‘bio-orthogonal’ and uniquely addressable 982 nt RNA scaffold sequence is folded by short DNA staple strands into a rectangle origami characterized by atomic force microscopy. In this context, the use of a split system, instead of a full aptamer sequence, conjugated to two RNA staple strands opens the possibility to monitor folding and unfolding, where multiple orthogonal split systems are considered, and to introduce other functionalization triggered by the assembly, providing a great potential for different applications.

We demonstrate the activation of the split system through the correct self-assembly of a well-defined rectangle shape (40 nm x 25 nm), able to fold isothermally at 53 °C within minutes. Finally, we demonstrate the application of the split Broccoli aptamer system for the sensitive *in vitro* detection of a specific DNA target sequence from the foodborne pathogen *Campylobacter* spp., mainly found in raw and undercooked meat. The two split sequences are conjugated into two distinct arms complementary to the *Campylobacter* spp. target sequence.^15^ Upon the addition of the single-stranded DNA target, the two non-functional RNA sequences hybridize to form the functional binding site, thus switching on the fluorescence signal of the specific dye. We designed and investigated three different systems as shortened variants, where the linker between the arm and the aptamer fragment consists of different nucleotide lengths. Then, the selected system was successfully tested by two optimized, simple and rapid assays: in-gel imaging and *in vitro* fluorescence measurements at 37 °C.

Looking forward, our results open the way to new sensitive aptameric genosensors and to protein-free tracking systems when long RNA scaffolds^16^ (e.g., messenger RNAs) or siRNA are delivered in living cells through therapeutics hybrid RNA/DNA nanostructures. Moreover, DNA-encoded RNA assemblies, circuits and memory structures^17^ carrying ‘light-up’ split aptamers can be designed, build and tested in cell-free systems as staging platforms before attempting engineering in living cells.

## RESULTS AND DISCUSSION

### Expression and self-assembly in an *E. coli* cell-free TX-TL system

Cell-free TX-TL systems and cell-free extracts are exciting tools that help characterise gene circuits and devices *in vitro* before moving *in vivo* for synthetic biology research: they are useful in gene network engineering, biomanufacturing and biosensing.^18-20^ Furthermore, cell-free systems offer an intermediate step to prototype nucleic acid-based materials^21^ and RNA assemblies, before moving the design to synthetic cells or more complex bacterial engineered cells.

The PURExpress system is one of the TX-TL platforms that combine bacteriophage T7 RNA polymerase and T7 promoter to the translational machinery of *E. coli*.^22^ In this paper, three components of a three-way junction carrying the split system were encoded in the corresponding dsDNA gBlocks containing the T7 promoter and T7 terminator (List S1). After amplification and purification, all the purified templates were transcribed, purified and checked by denaturing PAGE. As each transcript showed the expected size (Figure S1), the self-assembly at 37 °C of purified RNA sequences was at first demonstrated by in-gel imaging. The gel image (Figure S2) showed the selective DFHBI-1T staining of the three-way junction (lane 7) and the full Broccoli aptamer (positive control, lane 2), due to the aptameric G-quadruplex.

Computational prediction of the Broccoli three-way junction showed that the assembly is predicted even when T7 terminator sequences were included at the 3’ end of all RNA strands: the complex is formed when the concentrations of Split1d and Split2d are equal or greater than the concentration of the complementary strand (Figure S3). Furthermore, heatmaps underlined that Split1d and Split2d are not able to hybridise without the complementary sequence (Figure S3). Split1d/complementary and Split2d/complementary complexes exist as reported in Figure S3.

Finally, the three-way junction was expressed from double stranded templates and folded in the PURExpress system, enabling the fluorescence activation derived from the functional RNA Broccoli split system. As reported in Figure 1, after about 25 min at 37 °C the full assembly showed a fluorescence intensity of 1.84 (± 0.39, 3 independent measurements), while all the other samples (Split1d, Split2d, complementary sequence, Split1d or Split2d and complementary sequence, Split1d and Split2d) showed a fluorescence signal comparable to the basal fluorescence level of the cell-free system, underlined that the aptamer was not assembled. To further confirm the self-assembly of the three strands in the cells-free TX-TL system, aliquots were run on 10% TBE gel. As shown in Figure S4, the three-way junction assembled in the cell-free system (lanes 6 and 7, black arrows) is characterized by the same migration distance of the three-way junction assembled from transcribed and purified RNA (lane 3).

**Figure 1.**
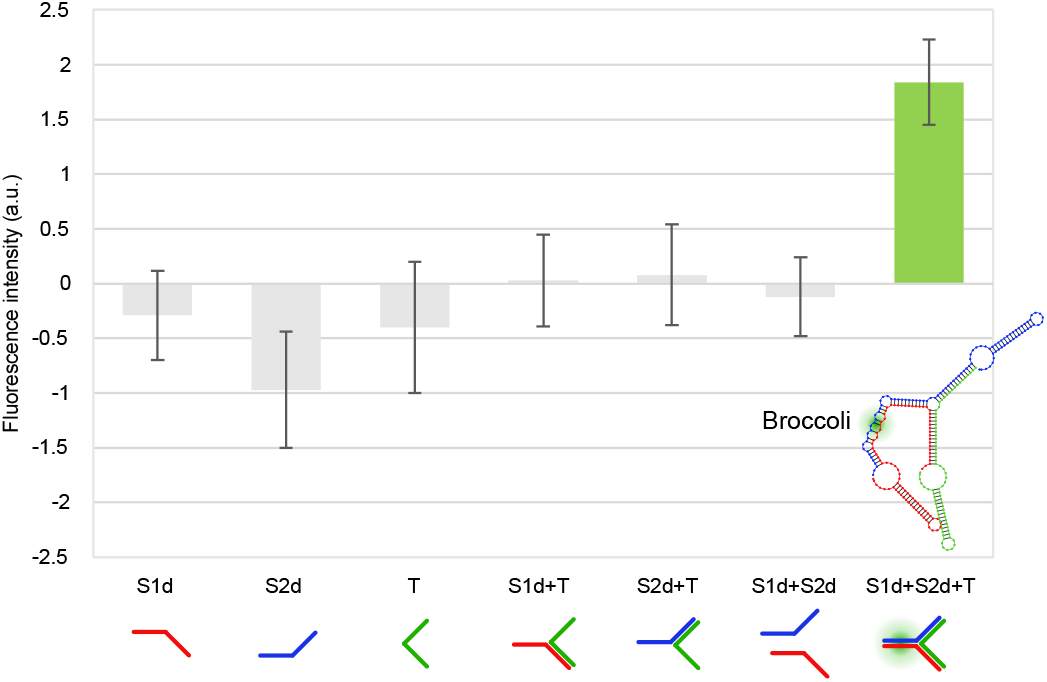
Fluorescence measurements in PURExpress cell-free TX-TL system containing gBlock templates. Fluorescence emission intensity is expressed in a.u. (F-F0, where F=sample fluorescence intensity; F0=cell-free system fluorescence without gBlock templates). From left to right: Split1d; Split2d; complementary sequence; Split1d and complementary sequence; Split2d and complementary sequence; Split1d and Split2d; Split1d, Split2d and complementary sequence (mean ± standard deviation, n=3).

### Activation of the Split Broccoli aptamer through a Hybrid RNA/DNA origami

Scaffolded DNA origami technology enables the bottom-up synthesis of nanostructures of defined shape and dimension, characterized by different functionalities and applications.^23^ In detail, a single-stranded DNA (ssDNA) scaffold sequence is folded into a precise geometry by several short ssDNA sequences, called staple strands.^24^

While many 2D and 3D DNA origami have been designed and visualized, hybrid RNA/DNA origami remains less explored despite the great potential to introduce functional RNA (e.g., RNA aptamers or siRNA) or mRNA as scaffolds.^25-29^

Recently, we proposed a custom-made software to select De Bruijn sequences (DBS of order 6) to be used as a scaffold in the origami synthesis. The obtained scaffolds are uniquely addressable and ‘bio-orthogonal’ by design and can be successfully folded into DNA or RNA origami^13,14^ and RNA/DNA hybrid origami.^27^

Here, we designed a hybrid ‘bio-orthogonal’ RNA/DNA origami rectangle (Figure 2) with dimensions of 40 nm x 25 nm using caDNAno.^30^ More specifically, the DBS scaffold had no repeats longer than 6 nt and excluded undesirable biological sequences, including start codon, Shine-Dalgarno sequence and restriction enzyme sites. A De Bruijn sequence of order *k*=6 has a sliding window of length 6 nt that is never repeated as it’s moved along the sequence.^27^

**Figure 2.**
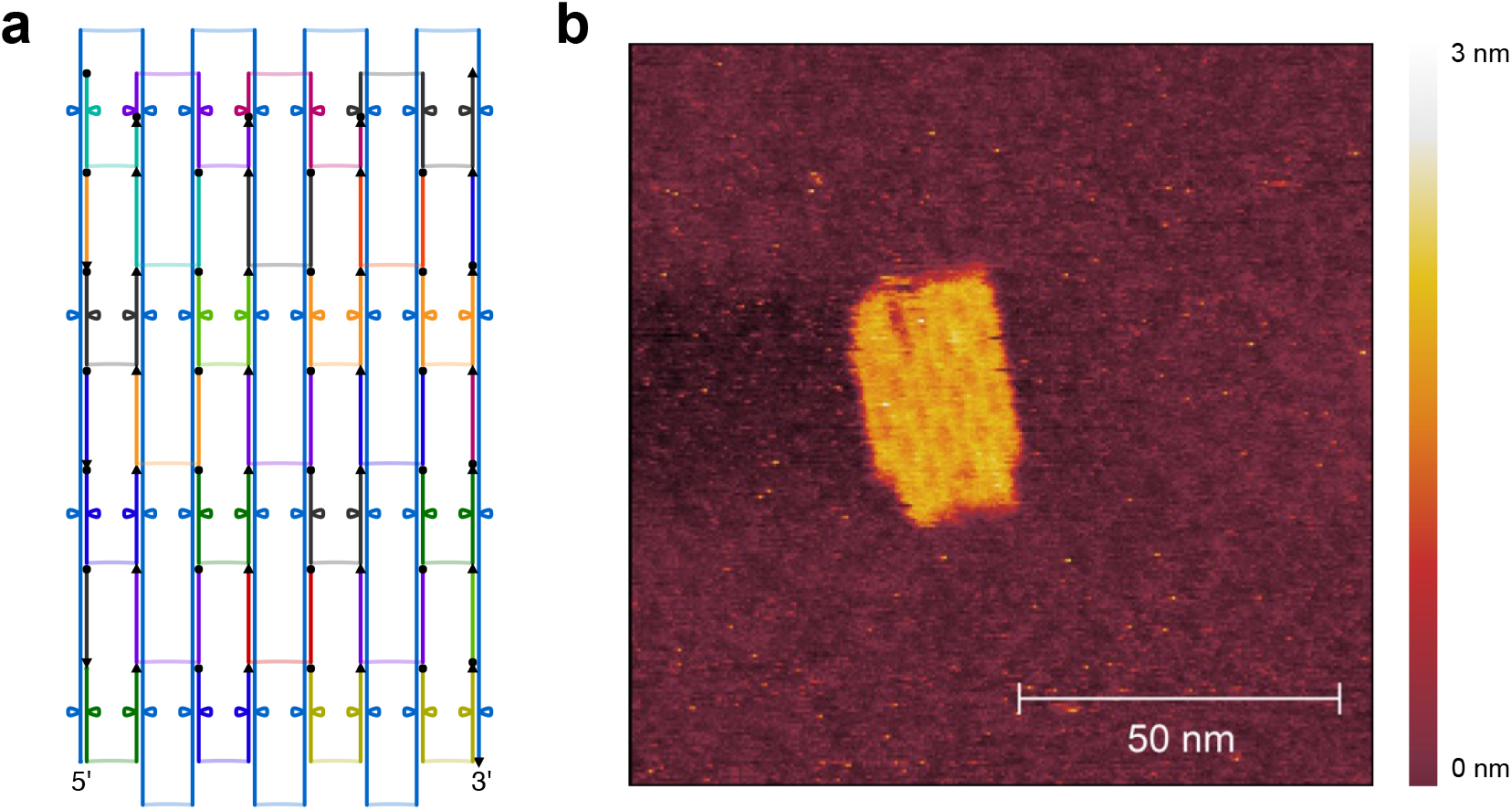
Scadnano schematic design (scaffold underlined in blue, staple strands shown in different colours, a) and high-resolution AFM image (b) of hybrid rectangle RNA/DNA origami.

The RNA scaffold (DBS 982) was synthesized by T7 transcription from a purified amplification product from plasmid pUCIDT-AMP-T7p-DBS982 (List S2): after denaturing PAGE, the purified transcript showed the expected size (Figure S5). The hybrid origami was then folded into a rectangle using 31 synthetic DNA staple strands (Table S1) and following a rapid isothermal protocol (53 °C for 15 min). To preliminary check the folding quality, the unpurified and purified RNA/DNA origami were run on 1.5% agarose gel. After electrophoresis, the gel image showed a distinct band for both purified and unpurified samples, suggesting proper self-assembly (Figure S6, lanes 4 and 5). The purified sample was visualized by AFM: the AFM images confirmed the correct formation of the origami with geometry and lengths consistent with the expected design (Figure 2, Figure S7 and Figure S8).

Finally, the split system was integrated into the hybrid origami characterized by AFM (Figure S9, green box). To demonstrate the functional activation of the split system through the origami self-assembly, samples were run in an agarose gel, stained with DFHBI, washed, and stained with SYBR® Gold. In a previous work,^31^ to verify the split malachite green aptamer activation through the self-assembly of an RNA cube characterized by cryo-EM, PAGE and fluorescence measurements were considered instead of a direct imaging. It should be noted that one dsDNA or aptamers protruding from 2D DNA origami were not well visible by AFM,^32,33^ probably due to the extension orientation with respect to the scanning direction.^33^

In our design, two DNA staples were converted to RNA sequences and elongated in 5’ and 3’ with Split1 and Split2. First, to demonstrate the fluorescence emission due to the hybridization between elongated staples S1, S2 and the complementary 36 nt short RNA derived from the DBS scaffold, we used specific DFHBI-1T in-gel imaging, a simple, selective and sensitive assay introduced by Jaffrey and coll.^8^ to monitor different expressed RNAs tagged with Broccoli. As shown in Figure S10, after the hybridization reaction and in-gel staining with DFHBI-1T, the 3-way junction assembly showed a clear fluorescent band (lane 7), thus demonstrating the ability of the system to turn on the fluorescence of the specific dye. Then, the split system was included in the hybrid origami: the purified origami sample was run on 1.5% agarose gel, with scaffold and staple strands as negative controls. After electrophoresis, the agarose gel was stained with DFHBI-1T and a fluorescent band related to the formation of the aptamer was imaged (Figure S11 left, lane 4, green arrow) at the migration distance corresponding to the purified hybrid origami (Figure S11 right, lane 4), while scaffold sequence and staple strands were unable to emit a fluorescent signal (Figure S12, lanes 4 and 5). It should be noted that compared to the previously described in-gel imaging using polyacrylamide gels, agarose gel staining with DFHBI-1T required a longer incubation time and higher PMT voltage (800 V) during laser scanning.

These results suggested that the split system can find potential application in studies related to *in vivo* delivery and tracking of hybrid origami using a non-cytotoxic dye. Furthermore, multiple split systems can be introduced in the same design with nanometer precision, increasing the DFHBI-1T emission signal.^31^

### Sequences design, dot blot assay on target CP3 and denaturing PAGE analysis of DNA/RNA oligonucleotides

To further confirm the versatility of our split system and to develop a proof-of-concept genosensor to *Campylobacter* spp., here we considered our split system and two analogues with a shorter 4 base pairs spacer arm or without a spacer. Indeed, as previously described, the choice of the split-site and the linkers between the affinity arms and the aptameric fragment are key factors to be considered when designing a binary aptamer for applications in diagnostics.^1,34^

In detail, the split sequences Split1 and Split2 with or without spacer arms (Split1/Split2-8nt, Split1/Split2-4nt and Split1/Split2-0nt, List S1) were conjugated in 5’ or 3’ end with RNA sequences complementary to a selected DNA target (CP3, 36 nt) from *Campylobacter* spp.^15,35,36^ (List S1), the most common bacterial cause of human gastroenteritis in the world.^37^ The 36 nt (18 nt on each split-sequence) oligonucleotide extension was designed on the 16S rRNA gene encoding for ribosomal *Campylobacter* RNA of *C. jejuni, C. coli, C. lari* and *C. upsaliensis*. Negative controls PE and PR corresponded respectively to a 36 nt *E. coli* sequence characterized by similarities to *Campylobacter*, and a 36 nt sequence designed by mismatching positions of the target sequence,^15^ resulting in a partial complementarity (12 bp) with Split1 sequence. The complementary biotinylated DNA detection probe (CampyP3) was used in a dot blot assay, as previously described:^15^ the obtained results showed a probe sensitivity of 1 ng/μL (Figure S13).

Since RNA and DNA sequences were used at specific molar concentrations and ratios, the purity of the designed split sequences, target CP3, negative controls PE and PR^15^ (List S1) was checked by denaturing PAGE (Figure S14, Figure S15 and Figure S16), loading ∼ 25 ng for each sample. The results showed RNA oligos, especially longer Split2 sequences, characterized by a lower purity compared to DNA sequences. As previously reported, the synthesis and purification of synthetic RNA oligonucleotides are less efficient compared to deoxynucleotides due to the 2’-OH protection, and the formation of several conformations (e.g., related to short intra-molecular double-stranded regions or stem-loop folding), especially for RNA longer than 50 nt.^38^ Nonetheless, considering the low micromolar concentrations (120 nM corresponding to ∼ 8 ng) to be used in the following in-gel imaging experiments, the RP-HPLC purified synthetic RNA oligos were not further purified.

Finally, the positive control Broccoli aptamer was transcribed from an amplified and purified dsDNA with a T7 promoter: the transcript was analysed by denaturing PAGE and the expected size was confirmed (66 nt, Figure S17).

### Split Broccoli aptamer system: PAGE analysis and in-gel imaging

To investigate the different split aptamer systems, Split1 and Split2 were mixed with CP3 (positive control), PE and PR (negative controls): after incubation at 37 °C for 25 min in the hybridization buffer supplemented with 10 mM MgCl_2_, samples were analysed by 10% PAGE. Split1/Split2-4nt, Split1/Split2-8nt and Split1/Split2-0nt showed an upper clear band, smeared and multiple bands or a faint band, respectively (Figure S18 b, a and c, lane 4). For these reason, Split1/Split2-4nt was selected to run the in-gel imaging experiment to confirm that the above mentioned upper band was the reconstituted aptamer upon CP3 hybridization (Figure 3 b). After electrophoresis, the 6% polyacrylamide gel was stained with 0.5 μM DFHBI-1T solution for 4 min. The gel imaging resulted in the selective staining of the hybrid assembly Split1/Split2-4nt/CP3 (Figure 3 c, lane 5, green arrow), while all the other loaded samples (Split1/Split2-4nt, CP3, PE, PR, Split1/Split2-4nt in the presence of PE or PR) were negative (Figure 3 c, lanes 3, 4, 6, 7, 8 and 9). It should be noted that higher DFHBI-1T concentration, incubation time and PMT voltage during the scanning, may result in undesirable non-specific staining. After fluorophore dye staining, the gel was washed three times and stained with SYBR® Gold to visualise RNA and DNA bands (Figure 3 d).

**Figure 3.**
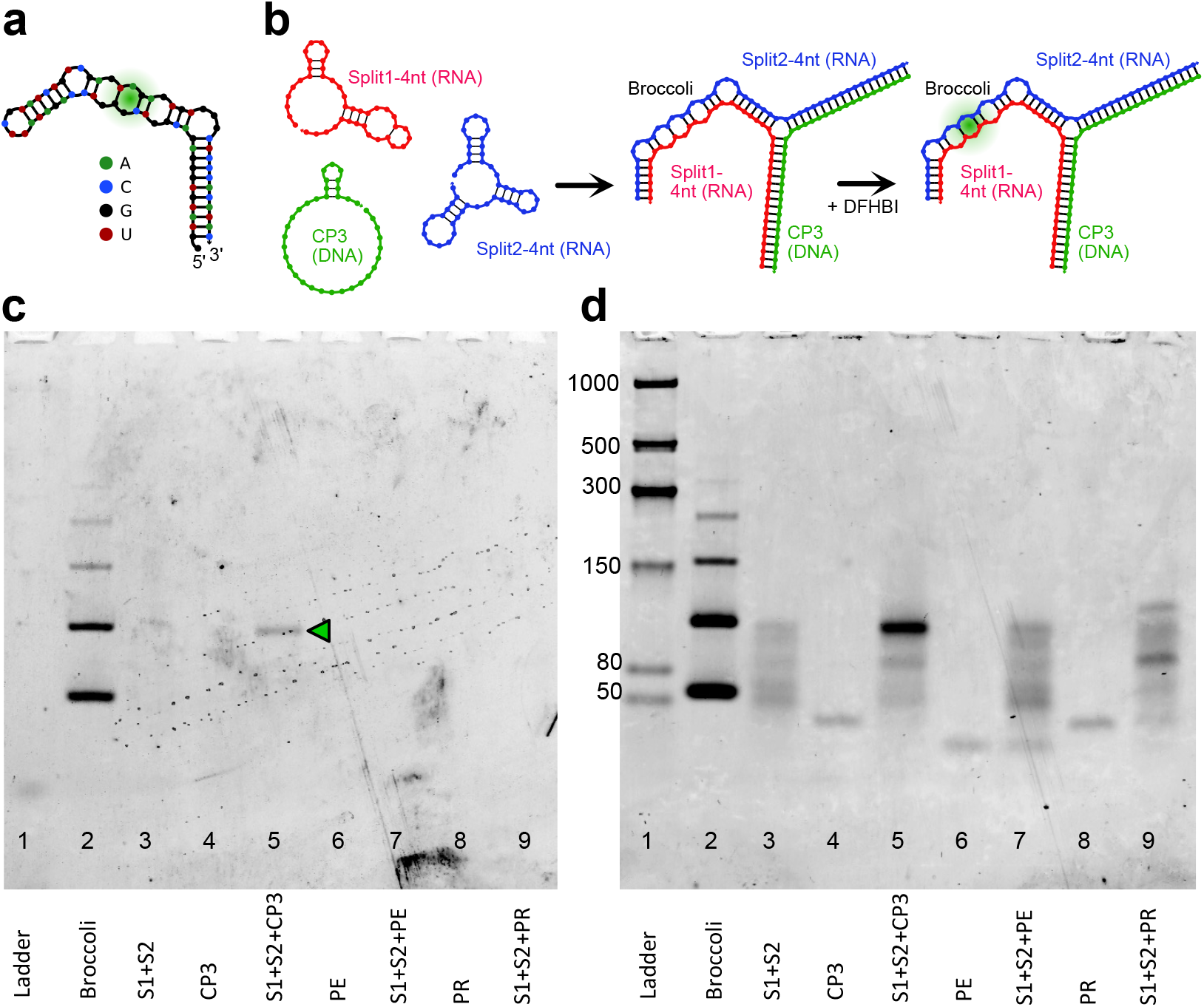
Split fluorescent Broccoli aptamer *in vitro* system. (a) Non-split Broccoli RNA aptamer sequence secondary structure. (b) Hybridisation of Split1-4nt, Split2-4nt and CP3 strands to yield a 3-strand complex with Broccoli aptamer. Addition of DFHBI causes Broccoli fluorescence. (c) and (d) In-gel imaging of the split system in the presence of CP3, PE or PR sequences. 6% TBE polyacrylamide gel after DFHBI-1T (c) and after SYBR® Gold (d) staining. The gel was stained with DFHBI-1T for 4 min to visualize Broccoli aptamer (positive control), and the hybridized Split1/Split2-4nt/CP3. After 3 washing steps, the gel was stained with SYBR® Gold for 5 min. Lanes: 1: low range ssRNA ladder; 2: Broccoli aptamer in (a); 3: Split1/Split2-4nt (0.12 μM each); 4: target CP3 (0.12 μM); 5: Split1/Split2-4nt and target CP3 (0.12 μM each); 6: PE (negative control, 0.12 μM); 7: Split1/Split2-4nt and PE (0.12 μM each); 8: PR (negative control, 0.12 μM); 9: Split1/Split2-4 nt and PR (0.12 μM each). Molecular sizes in nucleotides are indicated; the green arrow indicates the reconstituted split Broccoli aptamer in (b).

Finally, the double staining technique was applied to analyse samples with the same equimolar concentration of Split1/Split2-4 nt (0.12 μM) and reduced CP3 target concentration (0.10 μM, 0.08 μM and 0.05 μM). The lower detected CP3 concentration using DFHBI-1T corresponded to 0.6 ng/μL (Figure S19 a, lane 9). This result underlined that the simple in-gel imaging technique can be used as a rapid investigation tool when tagged hybrid DNA/RNA assembly are considered even at very low target concentration, thus demonstrating its sensitivity.

### Split Broccoli aptamer system: fluorescence measurements *in vitro*

The crystal structure of the Spinach-DFHBI complex revealed that potassium ions are part of the folded structure and stabilize the G-quadruplex, activating the fluorescence.^39^ As Broccoli aptamer retains the G-quadruplex of the DFHBI-binding site of Spinach aptamer, we postulated that Broccoli-DFHBI complex is characterized by a similar structure. Recently, we compared the fluorescence values of Broccoli aptamer in 20 mM Tris HCl pH 7.6, 1 mM EDTA, 10 mM MgCl_2_, and in 40 mM HEPES, 1 mM MgCl_2_, decreasing the potassium ions concentrations (from 100 Mm to 0 mM).^13^ Reducing K^+^ concentration in both buffers, resulted in a gradually lower fluorescence emission intensity with a higher emission at 100 mM and 50 mM KCl.^13^

The above described in-gel imaging protocol was performed in DFHBI-1T aptamer buffer (40 mM HEPES pH 7.4, 100 mM KCl, 1 mM MgCl_2_ and 0.5 μM DFHBI-1T), while the hybridization between the split system and target CP3 was conducted in the hybridization buffer containing 10 mM MgCl_2_, a typical magnesium ions concentration for the stability and folding of RNA in molecular biology experiments.^40^

Before moving from in-gel imaging to *in vitro* experiments, we first analysed by PAGE the hybridization products derived from reactions run in two different buffers: a) DFHBI-1T aptamer buffer without MgCl_2_ or containing 5 mM or 10 mM MgCl_2_; and b) hybridization buffer without KCl or supplemented with 50 mM or 100 mM KCl. Indeed, the split system performance as a genosensor should present a high emission fluorescence intensity signal and a low background, respectively correlated to DFHBI-complex stability (K^+^ dependent) and low Split1/Split2-4nt partial folding in the absence of the target CP3 (Mg^2+^ dependent). It should be noted that NUPACK analysis^41^ showed stable Split1-4nt and Split2-4nt secondary structures and a very low equilibrium concentration of Split1/Split2-4nt (0.00024 μM), considering the default RNA setting for salts (1 M Na^+^ and 0 M Mg^2+^), an incubation temperature of 37 °C and a concentration of 50 nM for each sequence (Figure S20 and Figure S21).

In the DFHBI aptamer buffer, the Split1/Split2-4 nt/CP3 hybrid band intensity increased considering the higher concentration of MgCl_2_, as expected (Figure S22 a, b and c, lane 4, black arrows). Overall, the band intensities were lower compared to the band intensities related to the same reaction conducted in the hybridization buffer (Figure S23 a, b and c, lane 4, black arrows). In buffers containing MgCl_2_ (5 mM or 10 mM), a faint band with the same migration distance appeared in samples Split1/Split2-4nt, Split1/Split2-4nt/PE and Split1/Split2-4nt/PR, presumably due to a low partial hybridization between Split1-4nt and Split2-4nt (Figure S22 b and c; Figure S23 a, b and c, blue arrows). Considering the results from PAGE analysis and to ensure a single predominant assembly in solution (Split1/Split2-4nt/CP3), the folding reactions and *in vitro* fluorescence measurements were initially performed in 20 mM Tris HCl pH 7.6, 1 mM EDTA, 100 mM KCl and 10 mM MgCl_2_ supplemented with DFHBI-1T.

After incubation at 37 °C for 15 cycles, the raw fluorescence values were measured by qPCR equipment. The Mg^2+^ concentration was reduced from 10 mM to 2.5 mM to further guarantee a weak hybridization between Split1 and Split2.

In the optimized buffer, Split1/Split2-4nt/PE and Split1/Split2-4nt/PR showed values closed to Split1/Split2-4nt, while Broccoli (17 ng) and Split1/Split2-4nt/CP3 (0.05 μM equimolar concentration) showed high fluorescence signals (Figure 4).

**Figure 4.**
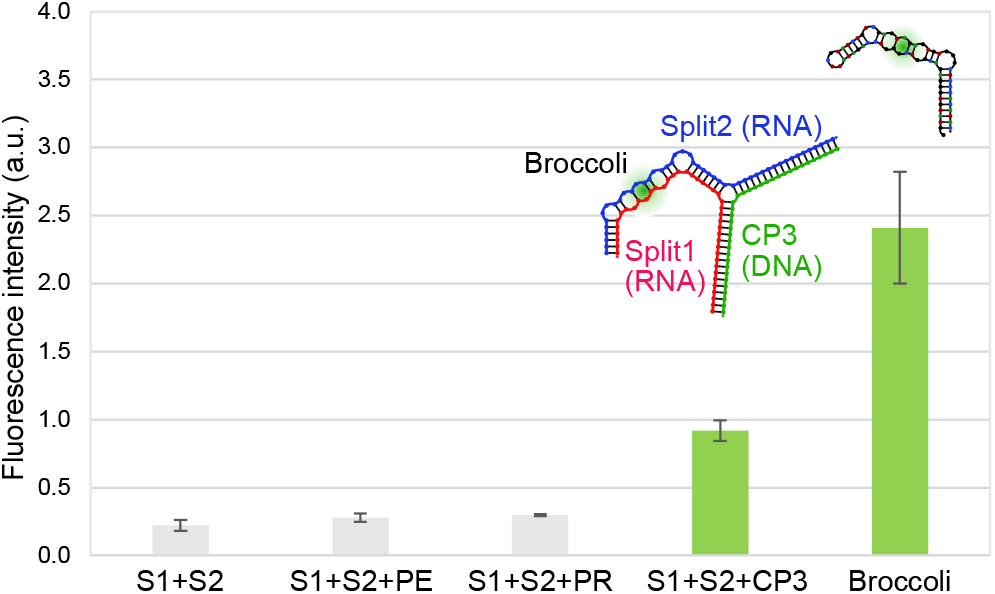
Fluorescence emission intensity in a.u. of the split system in the presence of equimolar concentration of PE, PR or CP3 sequences. From left to right: Split1/Split2-4 nt, Split1/Split2-4nt/PE, Split1/Split2-4nt/PR, Split1/Split2-4nt/CP3 (0.05 μM equimolar concentration) and Broccoli (17 ng), (raw data blank subtracted, mean ± standard deviation, n=3).

The sensitivity curve was obtained by subtracting the background fluorescence (Split1/Split2-4 nt) from the raw fluorescence data for different CP3 concentrations from 0.12 ng/μL to 0.57 ng/μL (3 independent measurements each, Figure 5). The correlation coefficient R^2^ was 0.995 and the limit of detection (LOD) was 0.074 ng/μL.

**Figure 5.**
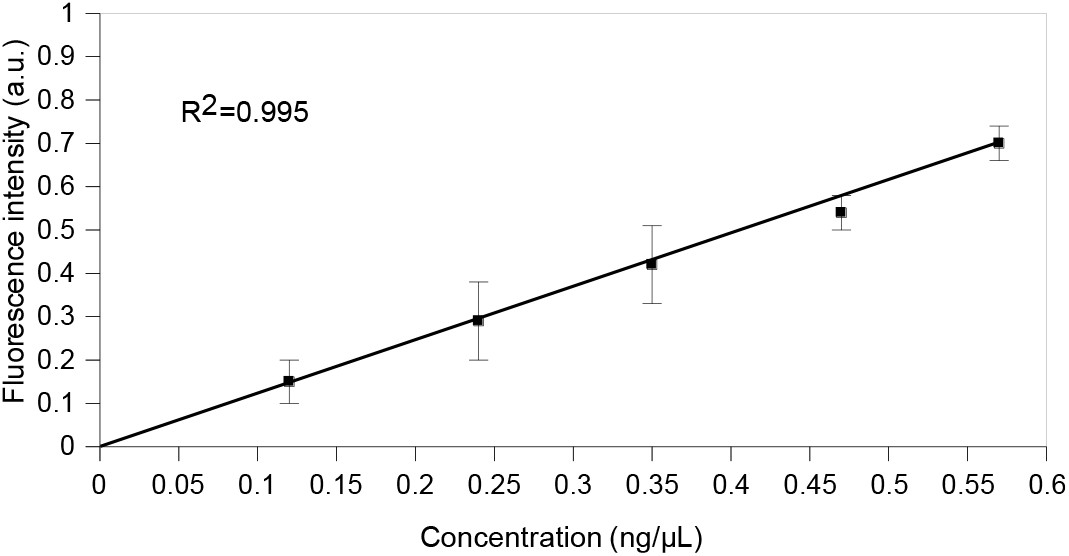
Sensitivity curve of the split system in the presence of different CP3 target concentrations. The fluorescence intensity in a.u. was obtained using Split1, Split2 (0.05 μM equimolar concentration) and different target CP3 concentration from 0.12 ng/μL to 0.57 ng/μL (0.01 μM, 0.02 μM, 0.03 μM, 0.04 μM and 0.05 μM), (raw data values are background subtracted; mean ± standard deviation, n=3).

## MATERIALS AND METHODS

### Reagents and Materials

All RNA oligonucleotides were purchased from Eurogentec and resuspended in Ultra Pure™ DNase/RNase free distilled water to give stock solutions of 100 μM and stored at – 80 °C. DNA oligonucleotides were purchased from Eurofins Genomics, Eurogentec or IDT. The pUCIDT-AMP-T7p_DB982 plasmid was purchased from IDT. The DNA oligonucleotides were resuspended in Ultra Pure™ DNase/RNase free distilled water to give stock solutions of 100 μM and stored at – 20 °C. The pUCIDT-AMP-T7p_DB982 plasmid was resuspended in 40 μL of IDTE buffer (10 mM Tris, 0.1 mM EDTA) and stored at – 20 °C. 0.5 M EDTA pH 8.0, Ultra Pure™ 10x Tris Borate EDTA (TBE) buffer, Ultra Pure™ 1 M Tris-HCl pH 7.5, Ultra Pure™ DNase/RNase free distilled water, 5 M NaCl (0.2 μm filtered), SYBR® Gold, 10x Tris Acetate EDTA (TAE) buffer molecular biology grade RNase free, 6% Novex™ TBE gel, 10% Novex™ TBE gel and 10% Novex™ TBE-Urea gel were purchased from Thermo Fischer Scientific. 1 M HEPES pH 7.0 Bioreagent, 1 M KCl BioUltra, 1 M MgCl_2_ BioUltra, nickel chloride, agarose and Nancy-520 were purchased from Sigma-Aldrich.

(5*Z*)-5[(3,5-Difluoro-4-hydroxyphenyl)methylene]-3,5-dihydro-2-methyl-3-(2,2,2-trifluoroethyl)-4*H*-imidazol-4-one (DFHBI-1T) was purchased from Tocris Bio-techne. A DFHBI-1T stock solution (20 mM) was prepared in DMSO (Sigma-Aldrich), stored in the dark at – 20°C and used within 4 weeks.

### Sequences design

Three RNA sequences^13^ were considered as components of a three-way junction and were cloned into dsDNA gBlocks containing T7 promoter and T7 terminator (Split1d, Split2d and complementary sequence): all the RNA sequences were analysed using NUPACK software^41^ and used in cell-free TX-TL experiments.

The uniquely addressable and bio-orthogonal synthetic RNA scaffold were generated with the computer code presented by Kozyra et al.^27^ The rectangle RNA-DNA hybrid origami (40 nm x 25 nm) was designed using caDNAno and scadnano software as described.^24,30,42,43^

To obtain the Split1/2-8nt system, two sequences complementary to the CP3 target were added at the 5’ and 3’ ends, and the 4-nt terminal stem-loop was removed as previously described.^13^ Split1/2-4nt and Split1/2-0nt showed a shorter or absent F30 arm, respectively. Split1 and Split2 are the two split Broccoli fragments. The RNA split sequences were analysed using NUPACK software^41^ the analysis was set at 37 °C, at a concentration of 50 mM for each RNA sequence and considering the salt concentration as 1 M Na^+^ and 0 M Mg^2+^ (default setting for RNA).

The target sequence CP3 (36 nt) was designed considering the 16S gene encoding for ribosomal *Campylobacter* RNA of *C. jejuni, C. coli, C. lari*, and *C. upsaliensis* (base location: 72–108), as previously reported.^15,35,36^ Non-complementary sequences (PR and PE, negative controls) were used to evaluate the selectivity of the Split1-2 systems. PR was designed by mismatching positions of the target sequence, while PE corresponded to a 36 nt *E. coli* sequence (accession number 527445.1, base location: 338–376) characterized by similarities to the *Campylobacter* sequence.^15^

### Expression, assembly and fluorescence measurements in PURExpress cell-free TX-TL system

The dsDNA gBlock templates (Broccoli, Split1d, Split2d and complementary sequence) containing the T7 promoter and terminator were amplified using the specific primers (0.25 μM or 0.5 μM) and Phusion® DNA polymerase (NEB). The amplification program was: 98 °C for 30 sec followed by denaturation at 98 °C for 10 sec, annealing for 15 sec and extension at 72 °C for 15 sec (15 cycles), additional extension for 5 min at 72 °C. The annealing temperatures were 68 °C (Broccoli), 67 °C (Split1) or 69 °C (Split2 and complementary sequence). After purification using Monarch® PCR & DNA Cleanup kit (NEB), the templates were transcribed: the purified transcripts were quantified and checked by NanoDrop One/OneC spectrophotometer and 10% TBE-Urea gel, respectively.

In order to verify the assembly products, RNA Split1d, Split2d and complementary (0.12 μM) were incubated at 37 °C for 25 min and then analysed by PAGE and in-gel imaging as described above. Cell-free experiments were performed using the PURExpress® system (NEB), following the manufacturer’s protocol. Briefly, 8 μL of solution A and 6 μL of solution B were mixed with 16 U RNase Inhibitor (NEB), gBlock templates (2 nM each) and 20 μM DFHBI-1T. The final reaction volume was adjusted to 20 μL with nuclease free water. All reactions (3 replicates for each combination) were set up on ice avoiding bubbles and incubated at 37 °C for about 25 min in a PCR cycler Rotor-Gene Q (Qiagen, excitation at 470 nm, emission at 515 nm, gain 10). It should be noted that special care is fundamental to avoid air bubbles, which are a source of variability and may interfere with fluorescence measurements increasing the possibility of outliers, a potential problem in cell-free experiments.^20^

Finally, 2.5 or 3.5 μL of cell-free reaction mix were incubated at 37 °C for about 50 min, run on 10% TBE gel at 200 V for 1 hour and 10 min and compared with the assembled purified transcript described above. All samples were stained with SYBR® Gold in 1x TBE for 25 min.

### Hybrid RNA/DNA origami synthesis and purification

A double-stranded DNA scaffold template was obtained from pUCIDT-AMP-T7p_DB982 plasmid (10 ng in 50 μL reaction mixture) by PCR amplification using Phusion® DNA polymerase (NEB), T7-DB982 forward and T7-DB982 reverse primers (0.5 μM each). An initial denaturation at 98 °C for 30 sec was followed by denaturation at 98 °C for 10 sec, annealing at 68 °C for 15 sec and extension at 72 °C for 15 sec (30 cycles). Finally, an additional extension was achieved for 10 min at 72 °C. The PCR product was purified using Monarch® PCR & DNA Cleanup kit (NEB) and eluted in 10 μL of Ultra Pure™ DNase/RNase free distilled water: the purified amplicon and the 1kb DNA ladder (NEB) were run on 1% agarose gel in 1x TBE for 1 h 30 min at 110 V. The gel was pre-stained with Nancy-520 and visualized under UV illumination (UVP GelStudio, AnalitikJena). The DNA concentration was measured on a NanoDrop One/OneC spectrophotometer (Thermo Scientific). The purified template was transcribed *in vitro* at 37 °C for 1 h using Ampliscribe™ T7-Flash™ Transcription kit (Lucigen, Epicentre) following the manufacturer’s instructions. After DNase treatment at 37 °C for 15 min, the RNA transcript was purified using RNA Clean & Concentrator™ (Zymo Research), eluted in 10 μL of Ultra Pure™ DNase/RNase free distilled water and quantified using a NanoDrop spectrophotometer. After the addition of the RNA loading dye 2x (NEB) and the heat denaturation at 65 °C for 5 min, *in vitro*-transcribed RNA scaffold was loaded and run on 10% TBE-Urea gel in TBE buffer at 200 V for 45 min. After staining with SYBR® Gold in 1x TBE for 5 min, the gel was visualized using Typhoon laser scanner. The low range ssRNA ladder (NEB) was used as molecular weight marker.

Single stranded DNA staple strands (final concentration 500 nM each) were mixed in a 50-fold excess with RNA scaffold in 50 μL of folding buffer (1x TAE molecular biology grade, 40 mM Tris-Acetate and 1 mM EDTA, supplemented with 12.5 mM MgCl_2_). The folding solution was incubated at 55 °C for 15 min and then held at constant temperature of 4 °C to stop the reaction. The folded constructs were purified from staple strands excess using the Amicon Ultra 0.5 mL 100 kDa centrifugal filters (Millipore). Capped Amicon Ultra were rinsed with 500 μL of RNase free folding buffer and centrifuged at 14000g for 1 min at 11 °C. After this preliminary washing step, the RNA/DNA origami samples were added to the filter device. Between every centrifugation step (14000g for 1 min at 11 °C repeated 5 times), the flowthrough was removed, and the filter was refilled with 450 μL of folding buffer. To recover the purified and concentrated origami sample, the filter was turned upside down and centrifuged at 1000g for 5 min at 11 °C. Scaffold, staple strands, unpurified and purified RNA/DNA origamis were run on 1.5% agarose gel in 1x TAE buffer at 90 V for 1 h at low temperature (below 10 °C). After staining with SYBR® Gold in 1x TAE for 8 min, the gels were visualized using a Typhoon laser scanner.

### Hybrid DNA/RNA origami characterization: Atomic Force Microscope (AFM) imaging

Freshly cleaved mica (Mica Grade V-4 12 mm Discs x 0.15 mm, Azpack Ltd) was passivated for 20 sec with 20 μL of 10 mM NiCl_2_ in Ultra Pure™ DNase/RNase free distilled water to ensure the adhesion of the negatively charged origami structures on the negative mica surface. After three washing steps with folding buffer (60 μL each), the purified origami solution was diluted 1:2 in the same buffer, added (10 μL) to the passivated mica surface, allowed to absorb for 1 min and imaged immediately (∼100 μL of the folding buffer were added after the origami absorption).

The purified hybrid RNA/DNA origami was characterized in tapping mode in liquid using VRS AFM Cypher ES (Asylum Research, Oxford Instruments, Santa Barbara, CA). The vertical oscillation of the BioLever Mini tip (spring k of 0.09 N/m, Asylum Research, Oxford Instruments, Santa Barbara, CA) was controlled by photothermal excitation (Blue Drive). All the images were lightly corrected using Gwiddion software.^44^

### Hybrid RNA/DNA origami and in-gel imaging

Two DNA staple strands were replaced with two protruding RNA staple strands including Broccoli split aptamer sequences. The hybridization reaction was run on 10% Novex™ TBE gel and analysed by in-gel imaging as previously described. Then, to confirm the incorporation of the Split Broccoli aptamer system into the purified rectangle nanostructure, the hybrid RNA/DNA origami was run on 1.5% agarose gel in TAE buffer (75 mL were poured into a 15 cm x 10 cm gel tray, Biorad) and analysed by in-gel imaging using the specific dye DFHBI-1T as described above with few modifications due to the different gel composition (agarose instead of polyacrylamide) and thickness. Briefly, the gel was washed three times for 5 min in RNase free water and then stained in a 12 cm square Petri dish for 1 h and 40 min with 50-60 mL of aptamer buffer containing 0.5 μM DFHBI-1T. The gel was imaged using Typhoon laser scanner (excitation 488 nm, emission 532 nm, normal sensitivity and PMT 800 V). Then, the gels were washed three times with RNase free water, stained with SYBR® Gold in 1x TAE for 10 min, and visualized using Typhoon laser scanner.

### Dot blot assay

The following samples were spotted onto the positively charged nylon membrane (Hybond™ XL, GE Healthcare): target CP3 (1 μL) at 100 ng/μL, 50 ng/μL, 10 ng/μL, 1 ng/μL, 0.1 ng/μL, 0.01 ng/μL, 1 pg/μL and 0.1 pg/μL, to check the sensitivity as previously described.^15^ Briefly, before deposition on membranes, DNA samples were denatured at 95 °C for 10 min and chilled immediately on ice. All samples (1 μL) were spotted on the nylon membranes and cross-linked to the air-dried membrane by exposure to UV light for 10 min. The membranes were soaked in a pre-warmed hybridization buffer at 65 °C for 30 min under gentle shaking. Hybridization was carried out at 65 °C overnight in the same buffer supplemented with 4 ng/μL of denaturated biotinylated CampyP3 probe (100 ng/μL). After incubation, the membranes were washed twice with 300 mM saline sodium citrate (SSC), 0.1% SDS for 5 min at room temperature on a shaker and then with 75 mM SSC for 15 min. The membranes were incubated in blocking solution at room temperature for 15 min and finally incubated with the same blocking solution supplemented with 0.7 μM streptavidin-HRP (30 min at room temperature). The signal was revealed using the enhanced chemiluminescent substrate for the detection of HRP (Thermo Fisher Scientific) under a ChemiDoc MP imaging system.^15^

### Synthesis of Broccoli aptamer and purity control of the split aptamer sequences (Split1,2-8nt, Split1,2-4nt and Split1,2-0nt) by denaturing PAGE

The double-stranded DNA template to be transcribed and containing the T7 promoter sequence was prepared by polymerase chain reaction (PCR) from Broccoli single-stranded DNA sequence (100 ng in 20 μL reaction mixture) amplified using Broccoli forward and reverse primers (0.5 μM) and Phusion® DNA polymerase (NEB). An initial denaturation at 98 °C for 30 sec was followed by denaturation at 98 °C for 10 sec, annealing at 64 °C for 20 sec and extension at 72 °C for 15 sec (20 cycles). Finally, an additional extension was achieved for 5 min at 72 °C. The PCR product was purified using Monarch® PCR & DNA Cleanup kit (NEB) and eluted in 10 μL of Ultra Pure™ DNase/RNase free distilled water: the purified amplicon and the low molecular weight DNA ladder (NEB) were run on 2% agarose gel in 1x TBE for 1 h 40 min at 100 V. The gel was pre-stained with Nancy-520 and visualized under UV illumination (UVP GelStudio, AnalitikJena). The DNA concentration was measured on a NanoDrop One/OneC spectrophotometer (Thermo Scientific).

The purified template was transcribed *in vitro* at 37 °C for 1 h and 30 min using Ampliscribe™ T7-Flash™ Transcription kit (Lucigen, Epicentre). After DNase treatment at 37 °C for 15 min, the RNA transcript was purified using RNA Clean & Concentrator™ (Zymo Research) and eluted in 10 μL of Ultra Pure™ DNase/RNase free distilled water and quantified using a NanoDrop spectrophotometer.

After the addition of the RNA loading dye 2x (NEB) and the heat denaturation at 65 °C for 5 min, *in vitro*-transcribed Broccoli aptamer was loaded and run on 10% TBE-Urea gel in TBE buffer at 200 V for 45 min. After staining with SYBR® Gold in 1x TBE for 5 min, the gel was visualized using Typhoon laser scanner and Image Quant TL software (normal sensitivity and PMT 380 V; GE Healthcare Life Sciences). The low range ssRNA ladder (NEB) was used as a molecular weight marker.

To check the synthesis purity, the chemically synthesized RNA and DNA sequences (split aptamer sequences, target CP3, non-complementary sequences PE and PR) were analysed by 10% TBE-Urea gel electrophoresis following the protocol described above.

### Assembly of RNA Split sequences with complementary CP3, non-complementary sequences PE and PR: PAGE and in-gel imaging

All the RNA and DNA sequences were diluted 1:10 from 100 μM stock solution. Each solution was quantified (ng/μL, average of 3 measurements) using NanoDrop One/OneC spectrophotometer: the exact molarity of each solution was obtained using software available online. RNA Split1 and Split2 sequences (8 nt, 4 nt or 0nt) were mixed in a 1:1:1 ratio (0.12 μM) with complementary CP3 or with negative controls PE or PR and incubated at 37 °C for 25 min in 20 mM Tris-HCl pH 7.5, 1 mM EDTA and 10 mM MgCl_2_ (hybridization buffer). To compare the hybridization of each Split1/Split2 system (8 nt, 4 nt or 0nt) with target and non-target sequences, all samples (6.5 μL) were run on 10% Novex™ TBE gel in 1x TBE buffer at 200 V for 40 min. After staining with SYBR® Gold in 1x TBE for 5 min, the gel was visualized using Typhoon laser scanner and Image Quant TL software (normal sensitivity and PMT 380 V, GE Healthcare Life Sciences). The low range ssRNA ladder (NEB) was used as a molecular weight marker.

For the in-gel imaging assay, Split1-4nt (0.12 μM) and Split2-4nt (0.12 μM) were mixed with target (0.12 μM, 0.10 μM, 0.08 μM and 0.05 μM of CP3) and non-target sequences (0.12 μM, PE or PR as negative controls). All samples (4.5 μL from 72 μL of reaction mix), including negative controls and Broccoli aptamer (positive control, 25-30 ng), were run on 6% Novex™ TBE gel in 1x TBE buffer at 180 V for 30 min. In-gel imaging was performed as previously described^8^ with some modifications. Briefly, the gels were washed three times for 5 min in RNase free water and then stained for 2-3 min in DFHBI aptamer buffer containing 40 mM HEPES pH 7.4, 100 mM KCl, 1 mM MgCl_2_ and 0.5 μM DFHBI-1T (50 mL of buffer in a 12 cm square Petri dish, Greiner Bio-One Ltd). The gels were imaged using a Typhoon laser scanner (excitation 488 nm, emission 532 nm, normal sensitivity and PMT 380 V). Then, the gels were washed three times with RNase free water, stained with SYBR® Gold in 1x TBE for 5 min, and visualized using a Typhoon laser scanner. The low range ssRNA ladder (NEB) was used as a molecular weight marker.

### Assembly of RNA Split sequences with complementary CP3, non-complementary sequences PE and PR: PAGE and fluorescence measurements by qPCR equipment

All the DNA and RNA sequences were prepared as described above. RNA Split1 and Split2 sequences (4 nt) were mixed in a 1:1:1 ratio (0.12 μM) with complentary CP3 or negative controls PE or PR and incubated at 37 °C for 25 min in different buffers. The used buffers were: a) 40 mM HEPES, 100 mM KCl and 1 mM MgCl_2_ (DFHBI-1T buffer), b) DFHBI-1T buffer supplemented with 5 MgCl_2_, c) DFHBI buffer supplemented with 10 MgCl_2_, d) 20 mM Tris-HCl pH 7.5, 1 mM EDTA and 10 mM MgCl_2_ (hybridization buffer), e) hybridization buffer supplemented with 50 mM KCl, f) hybridization buffer supplemented with 100 mM KCl. After the isothermal incubation, the resulting products were run on 10% Novex™ TBE gel in 1x TBE buffer at 200 V for 40 min. After staining with SYBR® Gold in 1x TBE for 5 min, the gel was visualized using Typhoon laser scanner and Image Quant TL software (normal sensitivity and PMT 380 V, GE Healthcare Life Sciences). The low range ssRNA ladder (NEB) was used as a molecular weight marker.

The fluorescence measurements in liquid were conducted using the real-time PCR cycler Rotor-Gene Q (Qiagen) considering the following protocol. Split1-4nt (0.05 μM) and Split2-4nt (0.05 μM) were mixed in ice with target (0.05 μM, 0.04 μM, 0.03 μM, 0.02 μM and 0.01 μM of CP3) and non-target sequences (0.05 μM, PE or PR as negative controls) in 20 mM Tris, 1 mM EDTA, 100 mM KCl and 2.5 mM MgCl_2_.

After the addition of DFHBI-1T solution (final concentration 0.5 μM) in ice, all samples (35 μL), including negative controls and Broccoli aptamer (positive control, 17 ng), were run in the real-time PCR cycler at 37 °C for 15 cycles (1 cycle/min) with excitation and emission at 470 nm and 515 nm, respectively (gain 10). The limit of detection (LOD) was calculated considering the standard deviation of the blank (10 ×3 independent measurements) and the calibration curve slope. All data were shown as mean ± standard deviation (SD) considering n=3. For each measurement, 3 readings were acquired and averaged.

## CONCLUSIONS

Here, we demonstrated the versatility of a Broccoli split aptamer system able to turn on the fluorescence signal of a non-cytotoxic and cell-permeable dye only in the presence of a specific RNA or DNA target sequence. At first, we showed that a designed three-way junction carrying the split system can be successfully expressed and assembled in a cell-free TX-TL system, opening the way to *in vivo* applications. Our results demonstrate the expression and assembly of a three-way junction restoring the functionality of the specific RNA ‘light-up’ aptamer in a reconstituted cellular environment. Accordingly, we underline the possibility to use a cell-free system as a technology to prototype functional RNA assemblies to be then expressed in synthetic cells or in bacterial cells. In this context, split fluorescent RNA aptamers can be used to detect specific hybridization or disassembly events as an alternative to Förster resonance energy transfer spectroscopy, unfeasible in cellular environment when expressing specific sequences.

As the recognition arms can be accurately changed, the split system was redesigned, and its functionality was successfully demonstrated through the self-assembly of a ‘bio-orthogonal’ hybrid RNA/DNA origami. We further confirm the versatility of our protein-free binary aptamer as a simpler reporter system compared to previously described approaches based on catalytic hairpin assembly circuit^10^ and *in situ* amplification method,^12^ in which the target is disconnected from the reporter system or may interfere with the cellular behaviour, respectively.

Finally, the RNA split aptamer sequences were elongated with a target recognition sequence specific for *Campylobacter* spp. After the selection of the Split1/Split2-4nt sequences and the optimization of the buffer composition, a sensitive genosensor was developed as a proof-of-concept for the specific DNA detection of femtomoles target at 37 °C in less than 20 minutes. This emphasizes that our split system could be used in specific DNA sequence detection and considered a promising tool for future biosensors development in alternative to molecular assays, such as real-time PCR with LOD of 10^3^ ng of DNA purified from meat samples.^45^ To further optimize our system by reducing the background noise, future developments can be focused on: i) immobilising a biotinylated split strand on streptavidin magnetic beads to introduce stringent washing steps normally used in dot blot assay; ii) adding destabilizing mismatches in the linker sequence of Split1-4nt and Split2-4nt, as previously mentioned by Kolpashchikov and coll.^1^

Because the reaction occurs isothermally at physiologically compatible temperature, the sensor can be genetically encoded, or the sensing reaction can be run where traditional diagnostic polymerase chain reaction is unfeasible.

Overall, our results may find applications in: i) DNA data storage structures^17^ labelling to monitor multiple hybridization steps or disassemblies *in vitro* or *in vivo*; ii) *in vivo* tracking of tagged-hybrid nanostructures used to deliver mRNA or siRNA as therapeutics; iii) new biosensing devices to detect specific DNA or RNA sequences.

## Authors contributions

Conceptualisation: N.K. and E.T. Methodology: E.T. and N.K. and M.M. Investigation: E.T., B.S.E., S.A.N., and P.V. Writing—original draft: E.T. Writing—review and editing: N.K., E.T., B.S.E., S.A.N., M.M., P.V. Supervision: N.K. and E.T. Project administration: N.K. Funding: N.K.

## Notes

The authors declare no competing financial interests.

## ACKNOWLEDGEMENTS

This work was supported by Engineering and Physical Sciences Research Council grant EP/N031962/1 and has received funding from the European Union’s Horizon 2020 research and innovation programme under grant agreement no. 899833. Krasnogor was supported by the Royal Academy of Engineering under the Chairs in Emerging Technologies scheme.

## ASSOCIATED CONTENT

### Supporting information

The Supporting information is available free of charge at

Details on the sequences, NUPACK prediction, dot blot assay, polyacrylamide gels, agarose gels, AFM images.

## Notes

### Competing Interest Statement

The authors have declared no competing interest.

