## Supplementary material for "Light-up split Broccoli aptamer as a versatile tool for RNA assembly monitoring in cell-free TX-TL system, hybrid RNA/DNA origami tagging and DNA biosensing": SUPPL INFO.docx

**List S1. dsDNA gBlock sequences and related primers, ssRNA split sequences, ssDNA CP3, PE and PR sequences (5'-3'; Split Broccoli sequences are underlined in green).**

**gBlock Broccoli**

TTCTAATACGACTCACTATAGGTATGTGGGAGACGGTCGGGTCCAGATATTCGTATCTGTCGAGTAGAGTGTGGGCTCCCACATACTAGCATAACCCCTTGGGGCCTCTAAACGGGTCTTGAGGGGTTTTTTG

**Forward primer gBlock Broccoli**

TTCTAATACGACTCACTATAGGTATGTGGG

**gBlock Split1d**

TTCTAATACGACTCACTATAGGTCCAAAAGAAGGAACCGTATGTGGGAGACGGTCGGGTCCAGATATAGCATAACCCCTTGGGGCCTCTAAACGGGTCTTGAGGGGTTTTTTG

**Forward primer gBlock Split1d**

TTCTAATACGACTCACTATAGGTCCAAAAGAAGGA

**gBlock Split2d**

TTCTAATACGACTCACTATAGGTATCTGTCGAGTAGAGTGTGGGCTCCCACATACTGGACGAAACTTCCCCTAGCATAACCCCTTGGGGCCTCTAAACGGGTCTTGAGGGGTTTTTTG

**Forward primer gBlock Split2d**

TTCTAATACGACTCACTATAGGTATCTGTCGAGT

**gBlock complementary target**

TTCTAATACGACTCACTATAGGGGAAGTTTCGTCCAGGTTCCTTCTTTTGGATAGCATAACCCCTTGGGGCCTCTAAACGGGTCTTGAGGGGTTTTTTG

**Forward primer gBlock complementary target**

TTCTAATACGACTCACTATAGGGGAAGTTTCG

**Reverse primer**

CAAAAAACCCCTCAAGACCCGTTTAG

**Split1-8nt**

UAGUGGCGCACGGGUGAGGUAUGUGGGAGACGGUCGGGUCCAGAUA

**Split2-8nt**

UAUCUGUCGAGUAGAGUGUGGGCUCCCACAUACUAAGGUAUAGUUAAUCUG

**Split1-4nt**

UAGUGGCGCACGGGUGAGGUGGGAGACGGUCGGGUCCAGAUA

**Split2-4nt**

UAUCUGUCGAGUAGAGUGUGGGCUCCCACUAAGGUAUAGUUAAUCUG

**Split1-0nt**

UAGUGGCGCACGGGUGAGGAGACGGUCGGGUCCAGAUA

**Split2-0nt**

UAUCUGUCGAGUAGAGUGUGGGCUCUAAGGUAUAGUUAAUCUG

***Campylobacter* P3** **target sequence (CP3)**

CAGATTAACTATACCTTACTCACCCGTGCGCCACTA

**Detection *Campylobacter* P3 sequence (CampyP3)**

[biotin] TAGTGGCGCACGGGTGAGTAAGGTATAGTTAATCTG

***E. coli* sequence EF527445.1 (PE)**

AAG ACG CGC ATC TCT TTT TTC ACC AGC GCG CTT TTC

**Random sequence (PR)**

CGT GCG CCA CTA CAG ATT ACC TTA AAC TAT CTC ACC

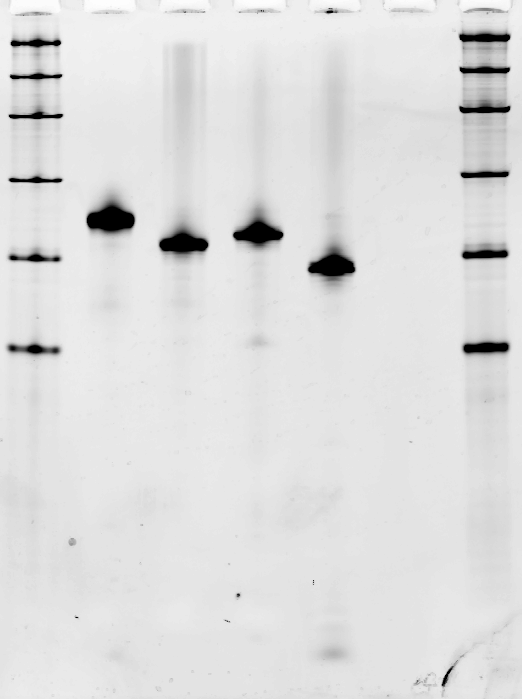

1	2	3	4	5	6

300

150

80

50

500

1000

**Figure S1. Broccoli aptamer and three-way junction sequences transcribed from double-stranded gBlock templates containing T7 promoter and T7 terminator.** 10% TBE-Urea gel electrophoresis after SYBR® Gold staining. Lanes. 1, 6: NEB low range ssRNA ladder; 2: Broccoli aptamer (113 nt); 3: Split1d (93 nt), 4: Split2d (98 nt); 5: complementary sequence (79 nt). All the sequences include the T7 terminator (47 nt). Molecular size in nucleotides are indicated.

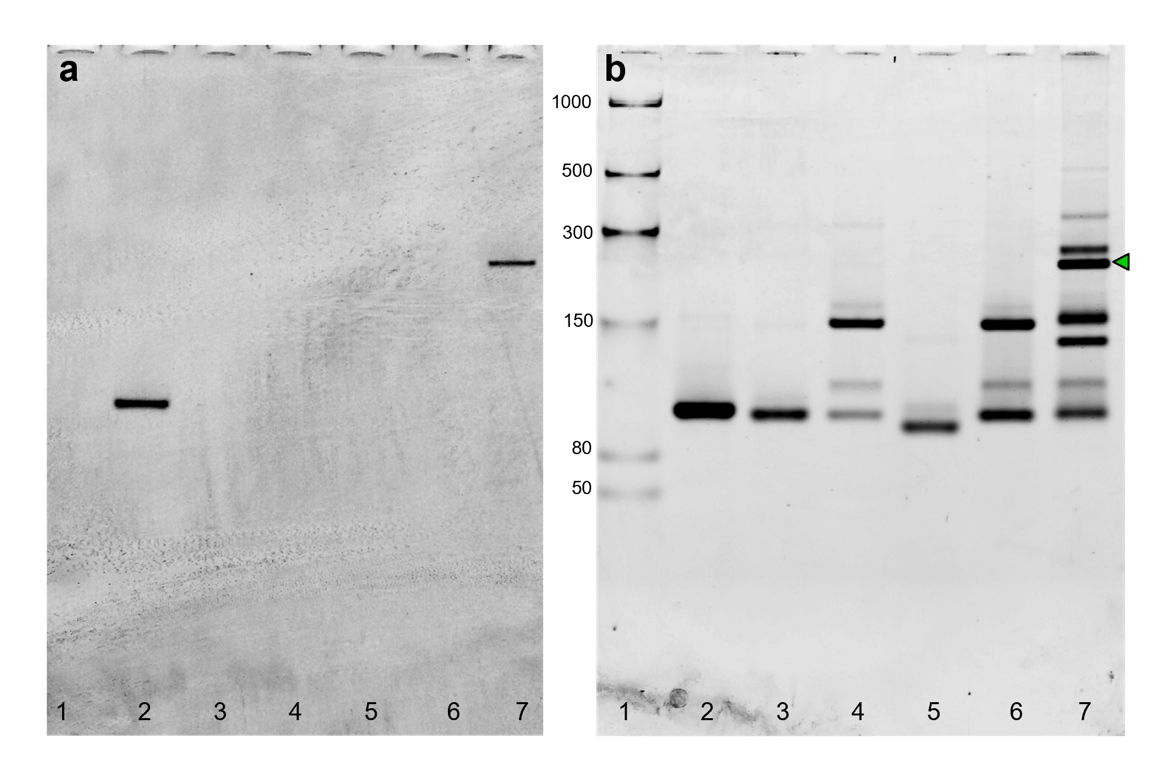

**Figure S2.** **In-gel imaging of Broccoli aptamer and three-way junction hybridised in vitro from transcribed sequences.** 6% TBE gel electrophoresis after DFHBI-1T (a) and after SYBR® Gold (b) staining. The gel was stained with DFHBI-1T for 3 min to visualize Broccoli aptamer (positive control), and the hybridized Split1d, Split2d and target. After 3 washing steps, the gel was stained with SYBR® Gold for 5 min. Lanes: 1: low range ssRNA ladder; 2: Broccoli aptamer; 3: Split1d; 4: Split2d; 5: target; 6: Split1d and Split2d; 7: Split1d, 2d and target. The green arrow underlines the 3-way junction assembly. Molecular size in nucleotides are indicated.

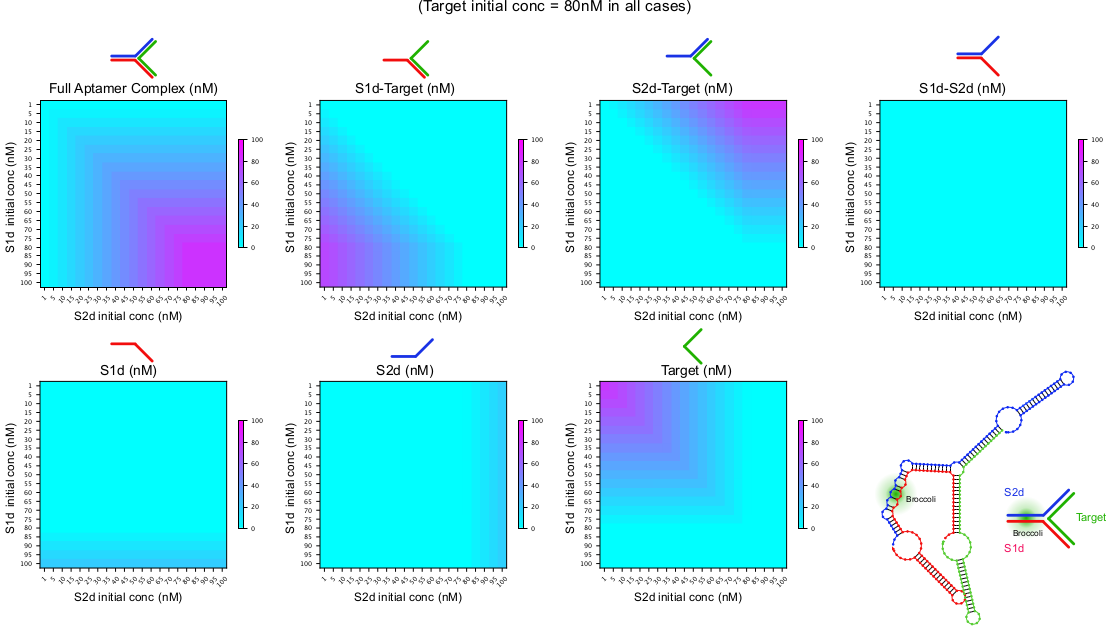

**Figure S3.** **Computational prediction of Broccoli 3-way junction complex formation under different *in vitro* concentration conditions.** The minimum free energy (MFE) structure of the full Broccoli aptamer complex is shown in bottom right. The MFE state is predicted to contain the Broccoli Aptamer region, even when T7 terminator sequences feature at the 3' end of all RNA strands. Heatmaps show the predicted equilibrium concentrations of the full Broccoli aptamer complex and other partial complexes as the initial concentrations of Split1d, Split2d are varied. The concentration of the target strand is constant at 80 nM in all plots. The heatmap’s results apply to the non/T7 terminator case and also to the T7 terminator case. From the heatmaps it can be observed that: (i) Split1d and Split2d are predicted not to hybridise without target (ii) Split1d-target complexes exist when Split2d is at relatively low concentration compared to Split1d and target (iii) Split2d-target complexes exist when Split1d is at relatively low concentration compared to Split2d and target (iv) The full 3-strand Broccoli complex is reliably formed when the concentrations of Split1d and Split2d are equal to, or greater than, the concentration of target. Figure created with NUPACK v4.0.0.27 test tube analysis.

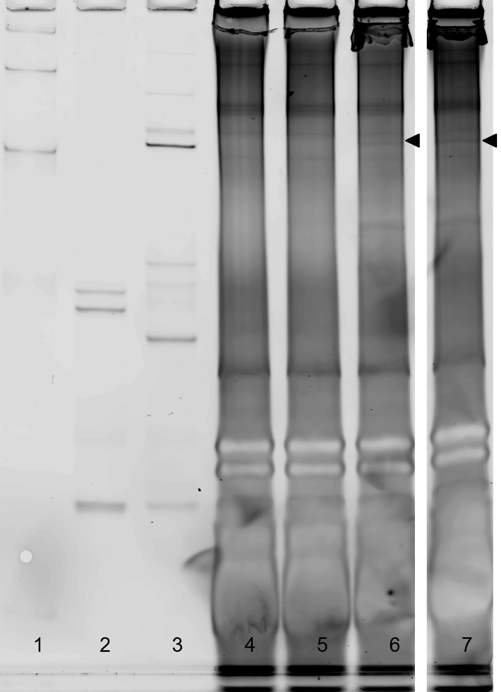

**Figure S4. PURExpress cell-free system containing the three-way junction gBlock templates analysed by polyacrylamide gel.** 10% TBE gel electrophoresis after SYBR® Gold staining. Lanes: 1: low range ssRNA ladder NEB; 2: Split1d and Split2d (after incubation at 37°C for 25 min); 3: Split1d, Split2d and target (after incubation at 37°C for 25 min); 4: cell-free system; 5: gBlock Split1d and Split2d in cell-free system; 6, 7: gBlock Split1d, Split2d and target in cell-free system loading 2.5 μL or 3.5 μL, respectively. Cell-free system samples (20 μL) are loaded after incubation at 37 °C for 50 min. The black arrow underlines the 3-way junction assembled in the cell-free system.

**List S2.** **De Bruijn scaffold sequence to be transcribed (inserted into the plasmid pUCIDT-AMP-T7p_DB982; in red T7 promoter sequence) and primers sequences (5'-3').**

TAATACGACTCACTATAGGCGGTCCAGCTAGCAGGTTTGCGGCTCAGAAGAGCTGTTGTGTTTGTTTTCGACTACCAGAACGGAGTCTCTAGCGTGAGATAAGTAAGATTAGGCTCGGAGAGTGTGAGGCTTCGTAATCGTACCACACACCAGGCGTAACCGCACTTAGACGCACAGGGTACAAGTGATAGGTAAAGTTACGGCAGGACGCCCAAAAGTCTGGAGCACAAACGGGGCCCCGCTAGGGAAAACGCCGGGGTAACTATTGTTATAATTCAAGAATTAGAACTAAAAGGTAGTAGCACCACTCGGTGGGTTAAACTAGCTAAAGACACCGCTCCAACAGCCGAAAGTGTACGCTGAATCACAGTCAAATTATACGGTGTTCGAGATCGCGAGTTTTGTGGGATTTGCACTCCAGATACCGATTCGGTAGCTTTATCGTTCACTGTGTCACGCGCAGCGCCACCAAAGCTGAGACGTTCTCGAAATTCTAATTTCTACGATTAAGTCCAAACAGAAAGCAATCTATTACACTGGAAGTCAGTAAAACAAAGGGATACAGATCCCGTGACGGCTAGTGCTGTGGTGTCCGAAGTTGACTGTCAGAGAACAATCGCACCGGACAGTTCGTTGAGTTCCAGTTGCAATTGCGATAGTAATAATAGATAGAGGCCGTGGAACCCCGTACTTCAGCGAGAAGTGGTCTTGGACTTGTACTGGGGCGAGCGGTGCGGGAACTCGTGTTGCCCGCAAGCACTGCAACACAGCGGAAGGATAGCAACGATCACTCTTGCTTTGTCGGACTCAGTCTAGGAGCCGCCGAGCCAGTCCCGCGCGTTCCCACGTTTCCGTAAACGTCCGCTTGGCCCGTCCACTGATATAGTTGGATCGGGAGAAATCGAAGCTCACGAACAGGAACGTAAGGCTGCTTGTTCTTTCACGGATCTCGGGCAGAATCTCAAACTCAATTACTCGATTTAGGTCGTCGCAGTACAG

**Primer Forward**

TAATACGACTCACTATAGGCGGTCC

**Primer Reverse**

CTGTACTGCGACGACCTAAATCG

**Table S1.** **Single-stranded DNA and RNA staple strands sequences (in green underlines the split Broccoli system).**

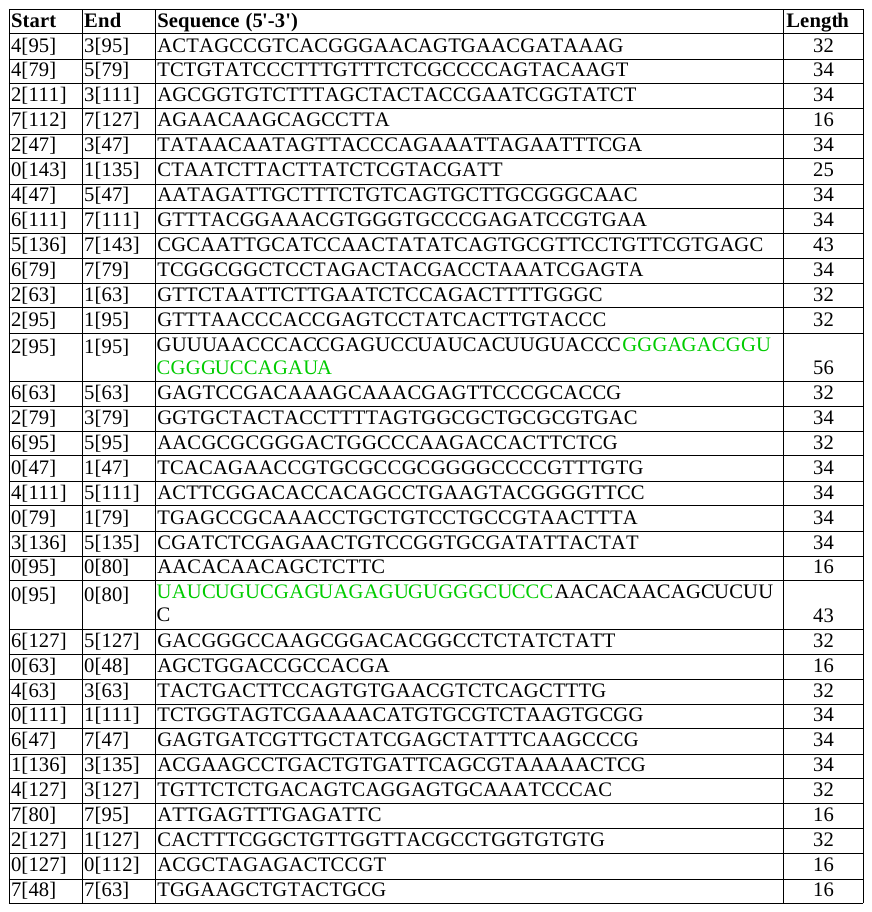

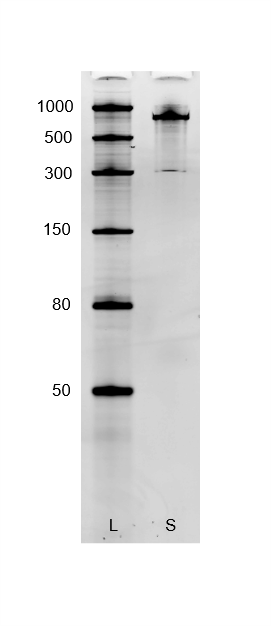

**Figure S5. Transcribed and purified De Bruijn scaffold sequence (DBS982) analysed by denaturing polyacrylamide gel.** 10% TBE-Urea gel electrophoresis after SYBR® Gold staining. Lanes. L: NEB low range ssRNA ladder; S: De Bruijn scaffold sequence (DBS982). Molecular sizes in nucleotides are indicated.

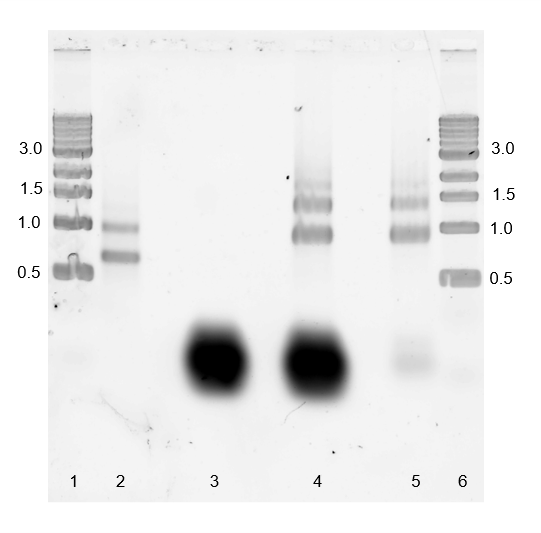

**Figure S6. Scaffold, staple strands, unpurified and purified hybrid RNA/DNA origami analysed by agarose gel.** 1.5% TAE agarose gel electrophoresis after SYBR® Gold staining. Lanes. 1 and 6: 1 kb DNA ladder; 2: transcribed and purified DBS982 RNA scaffold; 3: staple strands; 4: hybrid RNA/DNA origami; 5: hybrid RNA/DNA origami purified using Amicon Ultra centrifugal filter. Molecular sizes in kilobases are indicated.

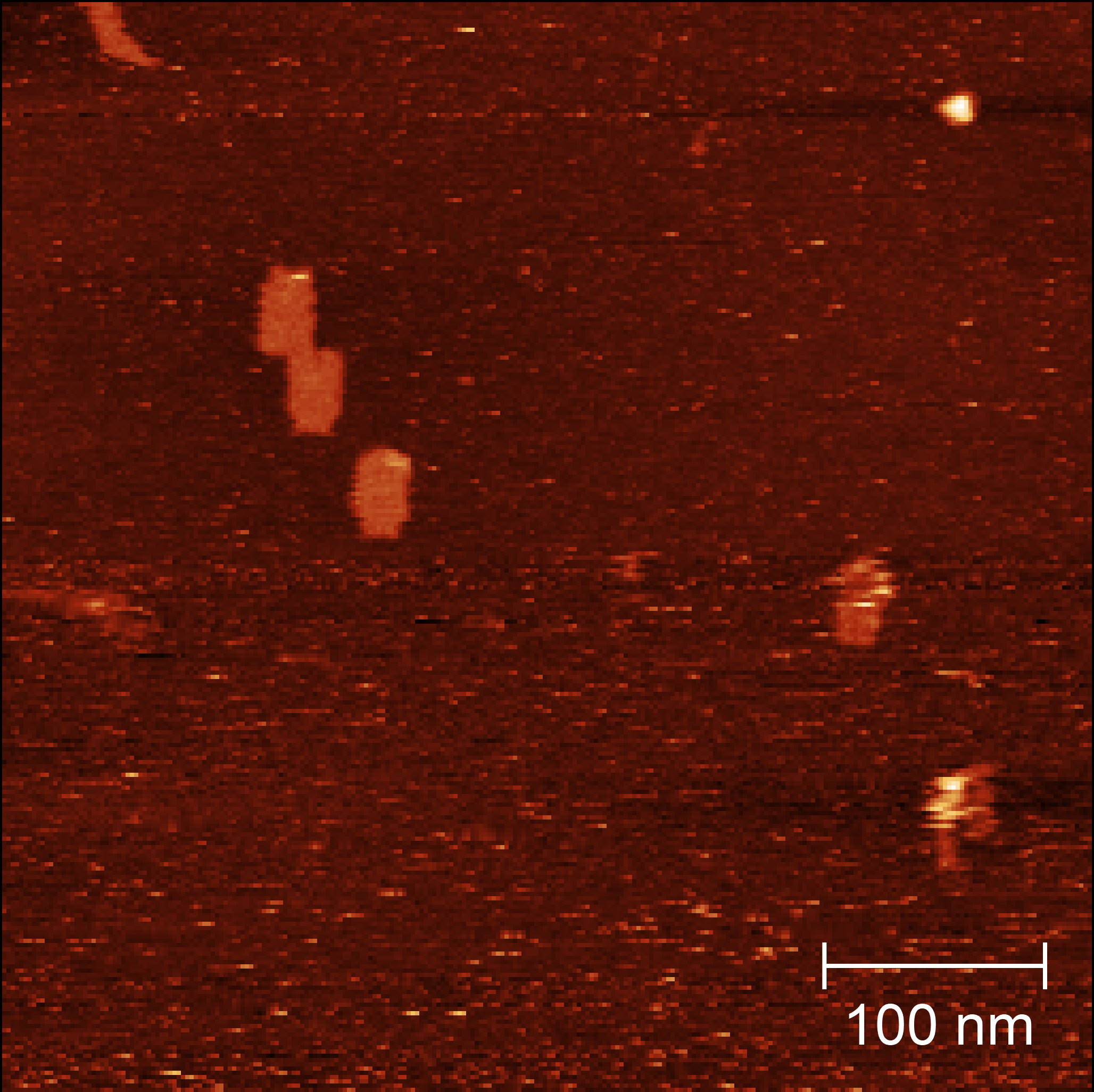

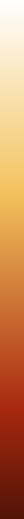

6 nm

0 nm

**Figure S7. High-resolution AFM image of hybrid rectangle RNA/DNA origami.**

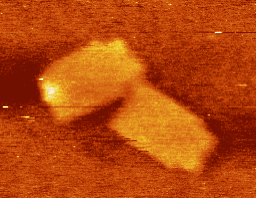

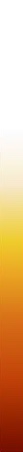

4 nm

0 nm

**Figure S8.** **High-resolution AFM image of hybrid rectangle DNA-RNA origami (scale bar 20 nm).**

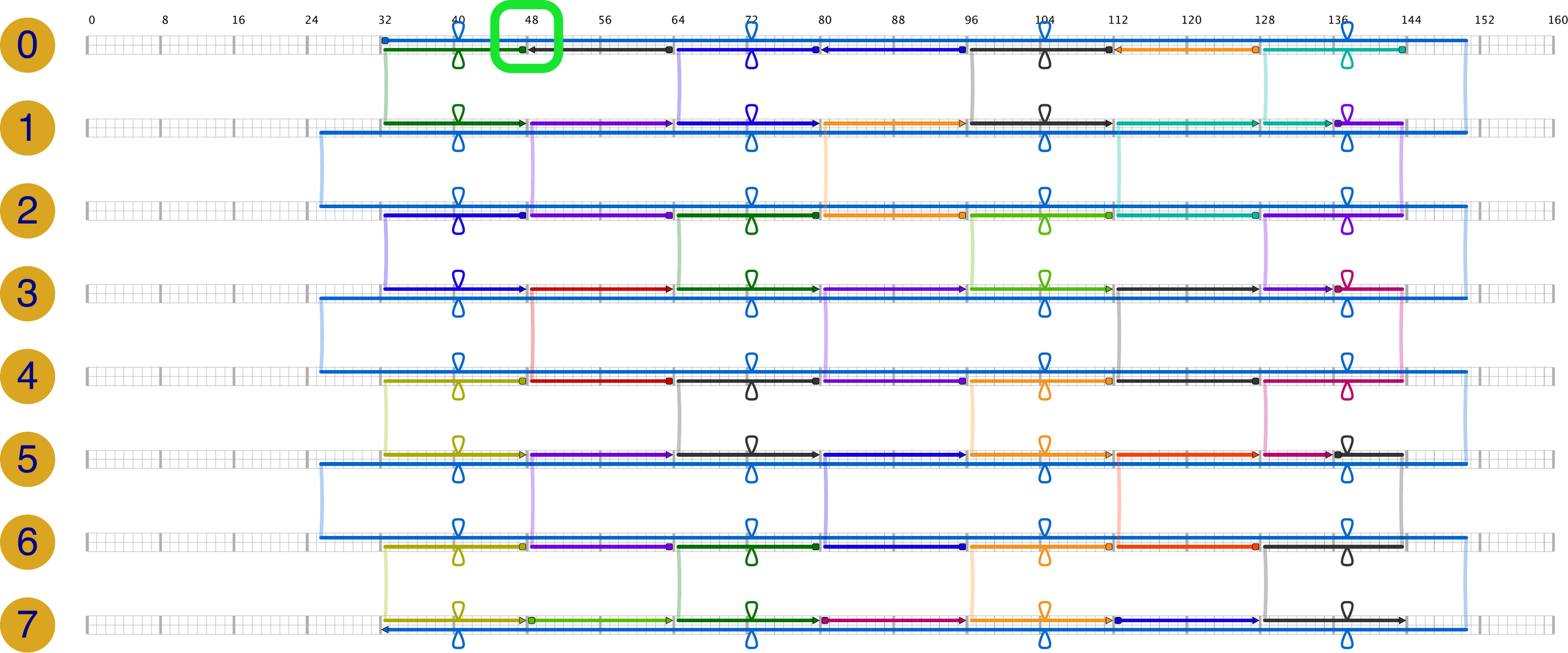

**Figure S9. Scadnano schematic design of the hybrid RNA/DNA origami.** The green box underlines the Split system position. The scaffold is underlined in blue, staple strands are shown in

different colours.

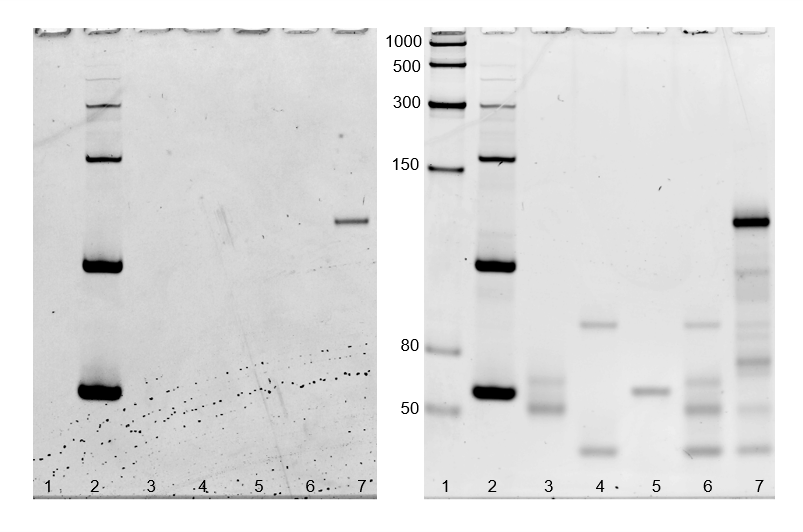

**Figure S10. In-gel imaging of staple S1 and staple S2 hybridized with complementary DBS sequence.** 10% TBE polyacrylamide gel after DFHBI-1T (left) and after SYBR® Gold (right) staining. The gel was stained with DFHBI-1T to visualize Broccoli aptamer (positive control), and the hybridized staple S1, staple S2 and complementary DBS target. After 3 washing steps, the gel was stained with SYBR® Gold for 5 min. Lanes: 1: low range ssRNA ladder; 2: Broccoli aptamer; 3: modified staple S1; 4: modified staple S2; 5: complementary DBS target sequence; 6: modified staple S1 and S2; 7: modified staple S1 and S2, and complementary DBS target sequence. Molecular sizes in nucleotides are indicated.

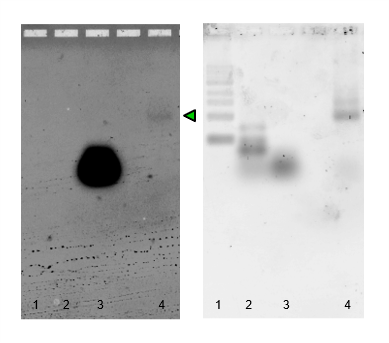

**Figure S11. In-gel imaging of Broccoli aptamer and purified RNA/DNA hybrid origami.** 1.5% TAE agarose gel after DFHBI-1T (left) and after SYBR® Gold (right) staining. The gel was stained with DFHBI-1T to visualize Broccoli aptamer (positive control), and the purified hybrid RNA/DNA origami. After 3 washing steps, the gel was stained with SYBR® Gold for 10 min. Lanes: 1: 1 Kb ladder; 2: low range ssRNA ladder; 3: Broccoli aptamer; 4: purified hybrid RNA/DNA origami (underlined by a green arrow).

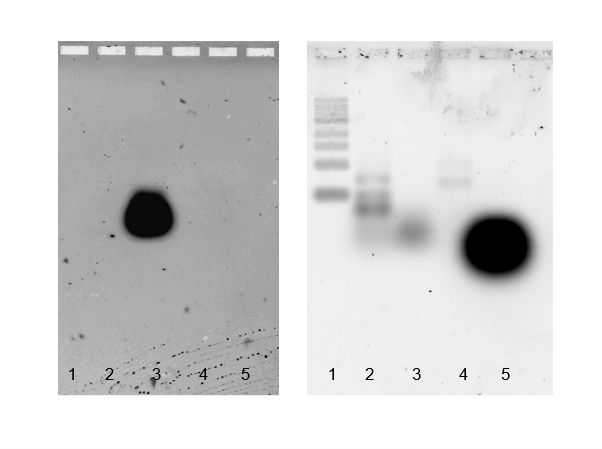

**Figure S12. In-gel imaging of Broccoli aptamer, RNA scaffold mix and staple strands mix.** 1.5% TAE agarose gel after DFHBI-1T (left) and after SYBR® Gold (right) staining. The gel was stained with DFHBI-1; after 3 washing steps, the gel was stained with SYBR® Gold for 10 min. Lanes: 1: 1 Kb ladder; 2: low range ssRNA ladder; 3: Broccoli aptamer; 4: RNA scaffold; 5: staple strands.

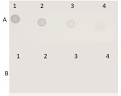

**Figure S13. Dot blot detection of biotin-labelled CampyP3 detection probe using different dilution of target complementary CP3 sequence. Row A.** A1: 100 ng/μL, A2: 50 ng/μL, A3: 10 ng/μL, A4: 1 ng/μL. **Row B.** B1: 0.1 ng/μL, B2: 0.01 ng/μL, B3: 0.001 ng/μL, B4: 0.0001 ng/μL.

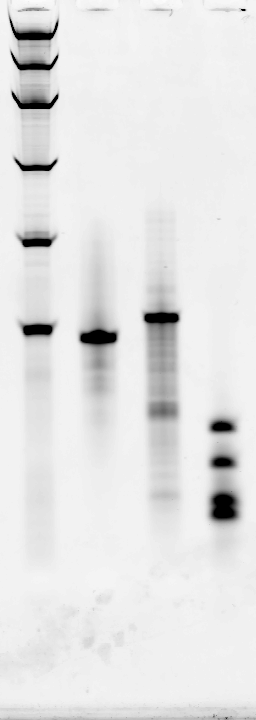

1000

300

150

80

50

29

25

21

17

500

1	2	3	4

**Figure S14. Split1-8nt and Split2-8 nt analysed by denaturing polyacrylamide gel.** 10% TBE-Urea polyacrylamide gel after SYBR® Gold staining. Lanes. 1: NEB low range ssRNA ladder; 2: Split1-8nt (46 nt), 3: Split2-8nt (51 nt); 4: ZR small-RNA^TM^ ladder. Molecular sizes in nucleotides are indicated.

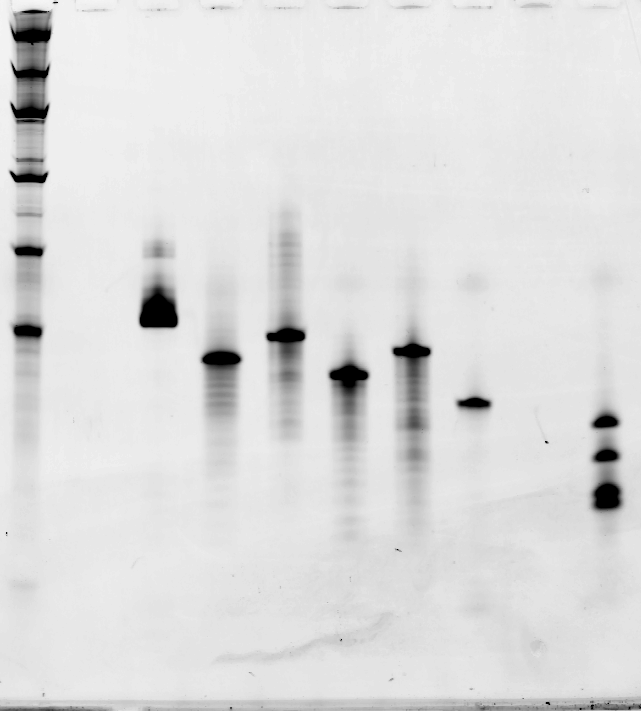

1000

300

150

80

50

1	2	3	4	5	6	7	8

29

25

21

17

500

**Figure S15. Broccoli aptamer, Split1-4nt, Split2-4nt, Split1-0nt, Split2-0nt and CP3 sequences analysed by denaturing polyacrylamide gel.** 10% TBE-Urea polyacrylamide gel after SYBR® Gold staining. Lanes. 1: NEB low range ssRNA ladder; 2: Broccoli aptamer (66 nt); 3: Split1-4nt (42 nt), 4: Split2-4nt (47 nt); 5: Split1-0nt (38 nt); 6: Split2-0nt (43 nt); 7: ssDNA CP3 target; 8: ZR small-RNA^TM^ ladder. Molecular sizes in nucleotides are indicated.

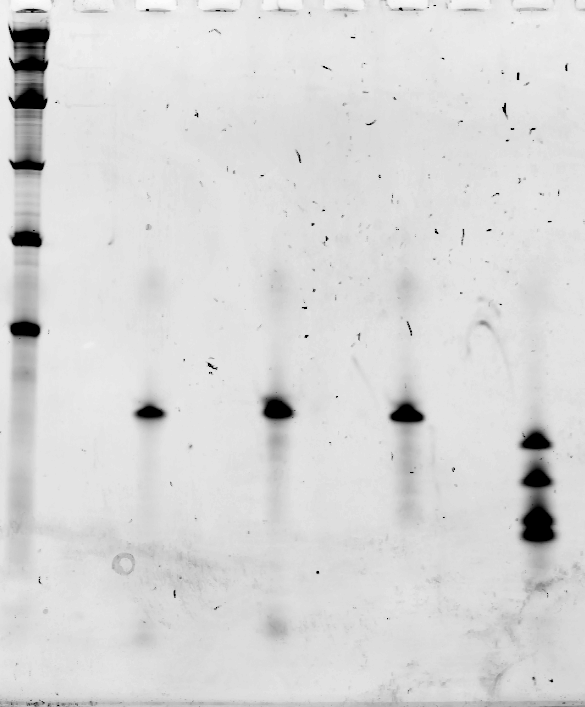

1000

300

150

80

50

29

25

21

17

1	2	3	4	5

500

**Figure S16. CP3, PE and PR sequences analysed by denaturing polyacrylamide gel.** 10% TBE-Urea polyacrylamide gel after SYBR® Gold staining. Lanes. 1: low range ssRNA ladder; 2: ssDNA target CP3; 3: ssDNA PE (negative control); 4: ssDNA PR (negative control); 5: ZR small-RNATM ladder. Molecular sizes in nucleotides are indicated.

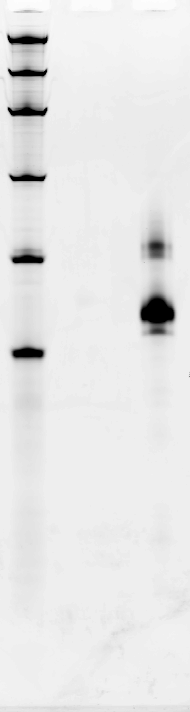

L	B

1000

300

150

80

50

500

**Figure S17. Transcribed and purified broccoli sequence analysed by denaturing polyacrylamide gel.** 10% TBE-Urea polyacrylamide gel after SYBR® Gold staining. Lanes. L: low range ssRNA ladder; B: Broccoli aptamer. Molecular sizes in nucleotides are indicated.

1000

300

150

80

50

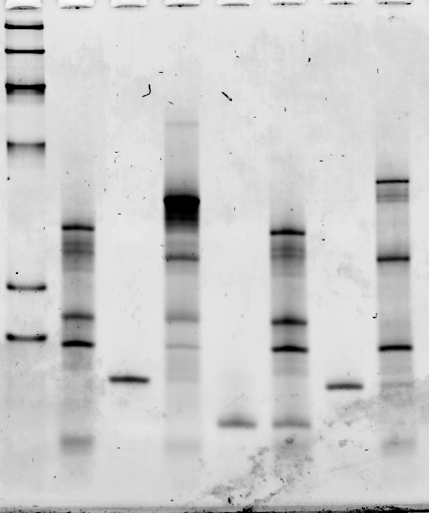

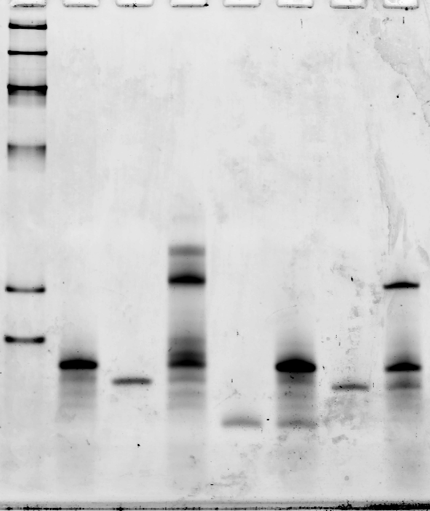

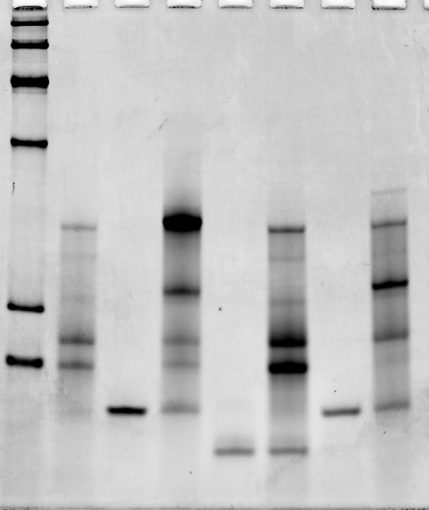

1	2	3	4	5	6	7	8

**b**

**c**

**a**

1	2	3	4	5	6	7	8

1	2	3	4	5	6	7	8

**Figure S18. Comparison of hybridization reactions in 20 mM Tris-HCl pH 7.5, 1 mM EDTA and 10 mM MgCl_2,_ using Split1/Split2-8nt, Split1/Split2-4nt or Split1/Split2-0nt and CP3, PE or PR sequences.** 10% TBE polyacrylamide gel after SYBR® Gold staining. **a**: Split1/Split2-8nt; **b**: Split1/Split2-4nt; **c**: Split1/Split2-0nt. Lanes: 1: low range ssRNA ladder; 2: Split1 and Split2; 3: target CP3; 4: Split1/Split2 and target CP3; 5: PE; 6: Split1/Split2 and PE (negative control); 7: PR (negative control); 8: Split1/Split2 and PR. Molecular sizes in nucleotides are indicated.

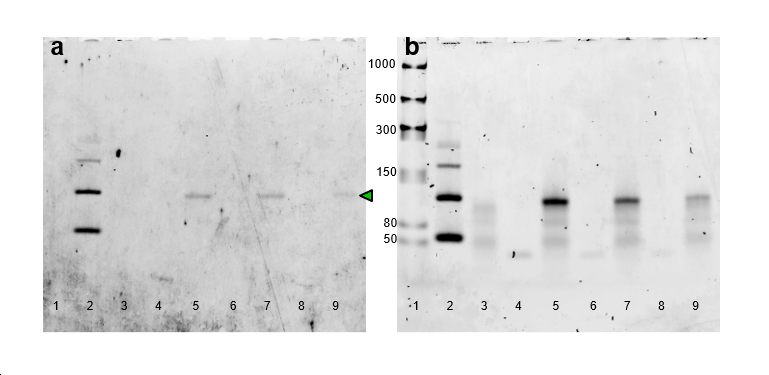

**Figure S19. In-gel imaging of the split system in the presence of CP3 target sequences at different concentrations.** 6% TBE polyacrylamide gel after DFHBI-1T (a) and after SYBR® Gold (b) staining. The gel was stained with DFHBI-1T for 4 min to visualize Broccoli aptamer (positive control), and the hybridized Split1/Split2-4nt/CP3. After 3 washing steps, the gel was stained with SYBR® Gold for 5 min. Lanes: 1: low range ssRNA ladder; 2: Broccoli aptamer; 3: Split1/Split2-4nt (0.12 µM); 4: target CP3 (0.10 µM); 5: Split1/Split2-4nt (0.12 µM each) and target CP3 (0.10 µM); 6: target CP3 (0.08 µM); 7: Split1/Split2-4nt (0.12 µM each) and target CP3 (0.08 µM); 8: target CP3 (0.05 µM); 9: Split1/Split2-4nt (0.12 µM each) and target CP3 (0.05 µM). Molecular sizes in nucleotides are indicated; the green arrow underlines the reconstituted split Broccoli
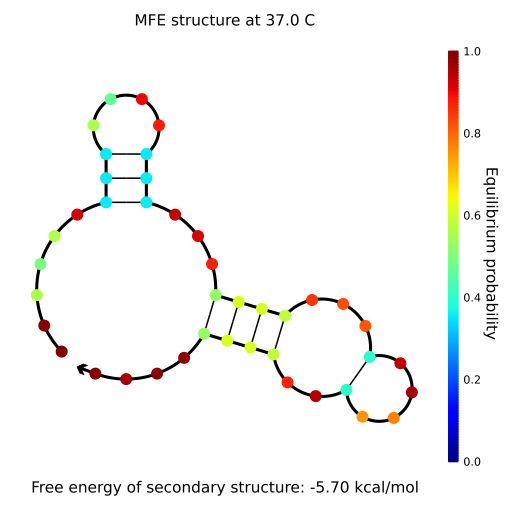
aptamer.

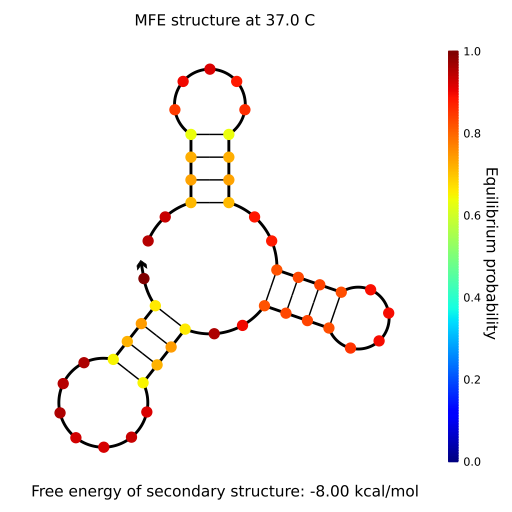
**Figure S20. NUPACK secondary structure prediction of Split1-4nt sequence (50 nM) at 37 °C in 1 M Na^+^ and 0 M Mg^2+^.**

**Figure S21.** **NUPACK secondary structure prediction of Split2-4nt sequence (50 nM) at 37 °C in 1 M Na^+^ and 0 M Mg^2+^.**

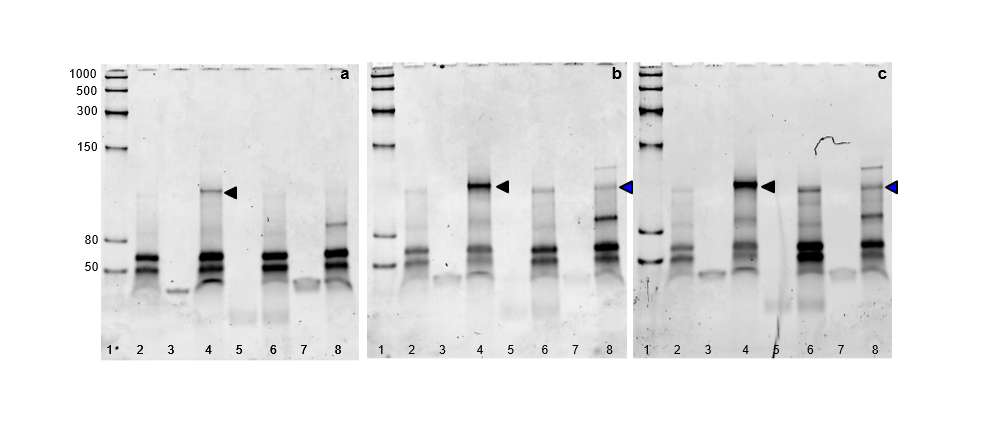

**Figure S22. Comparison of hybridization reactions using Split1/Split2-4nt and CP3, PE or PR sequences in DFHBI-1T aptamer buffer without or with different MgCl_2_ concentrations.** 10% TBE polyacrylamide gel after SYBR® Gold staining. Hybridization reactions using Split1/Split2-4nt are performed in: a) DFHBI-1T aptamer buffer; b) DFHBI-1T aptamer buffer supplemented with 5 mM MgCl_2_; c) DFHBI-1T aptamer buffer supplemented with 10 mM MgCl_2_. Lanes: 1: low range ssRNA ladder; 2: Split1-4nt and Split2-4nt; 3: target CP3; 4: Split1/Split2-4nt and target CP3; 5: PE; 6: Split1/Split2-4nt and PE (negative control); 7: PR (negative control); 8: Split1/Split2-4nt and PR. Molecular sizes in nucleotides are indicated. Black arrows: Split1/Split2-4nt/CP3 hybrid; blue arrows: low partial hybridization between Split1-4nt and Split2-4nt.

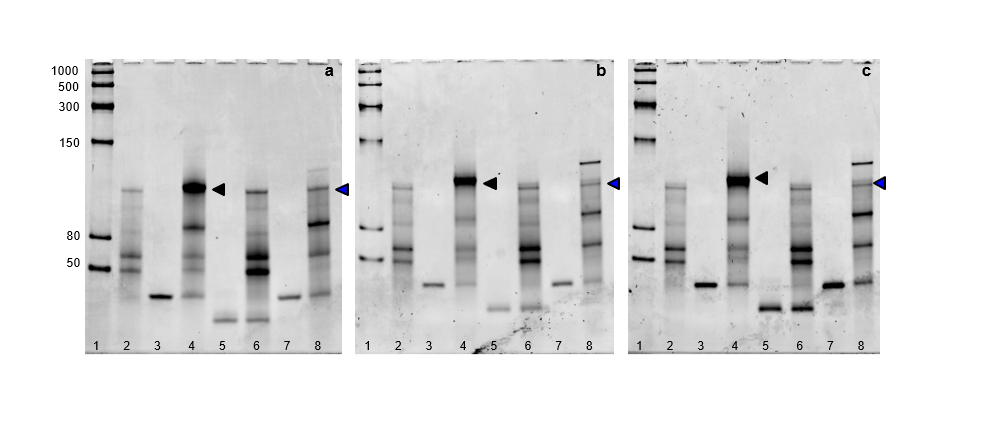

**Figure S23. Comparison of hybridization reactions using Split1/Split2-4nt and CP3, PE or PR sequences in hybridization buffer without or with different KCl concentrations.** 10% TBE polyacrylamide gel after SYBR® Gold staining. Hybridization reactions are performed in: a) 20 mM Tris HCl pH 7.6, 1 mM EDTA, 10 mM MgCl_2_ (hybridization buffer); b) 20 mM Tris HCl pH 7.6, 1 mM EDTA, 10 mM MgCl_2_ supplemented with 50 mM KCl; c) 20 mM Tris HCl pH 7.6, 1 mM EDTA, 10 mM MgCl_2_ supplemented with 100 mM KCl. Lanes: 1: low range ssRNA ladder; 2: Split1-4nt and Split2-4nt; 3: target CP3; 4: Split1/Split2-4nt and target CP3; 5: PE; 6: Split1/Split2-4nt and PE (negative control); 7: PR (negative control); 8: Split1/Split2-4nt and PR. Molecular sizes in nucleotides are indicated. Black arrows: Split1/Split2-4nt/CP3 hybrid; blue arrows: low partial hybridization between Split1-4nt and Split2-4nt.
